# Chemical probes reveal individualized gut microbiome biotransformation capacity and the impacts of ex vivo fermentation conditions

**DOI:** 10.64898/2026.07.30.741225

**Authors:** Jacob Folz, Raúl Fernández Cereijo, Maja Stevanoska, Hélène Bigonne, Georg Aichinger, Shana Sturla

**Affiliations:** ETH Zurich, Laboratory of Toxicology

## Abstract

The gut microbiome transforms endogenous and exogenous chemicals, contributing to bioactivation or detoxification via the formation of metabolites with altered bioactivity. Most high content microbiome assays infer function from genetic composition rather than direct assessment of biotransformation activity, and it remains difficult to predict functional consequences of environmental factors and experimental variation. Therefore, we developed an anaerobic fecal fermentation workflow that couples targeted LC-MS/MS quantification of dynamic profiles of 20 chemical probes with untargeted metabolomics to profile human microbiome biotransformation capacity and used it to assess the impact of experimental conditions on biotransformation profiles. Across five donors and 240 fermentations, inoculum density and growth medium composition strongly influenced probe transformation rates, whereas the biotransformation capacities of fecal slurries frozen at −80°C did not differ from fresh fecal samples. Individual donors could be uniquely stratified on the basis of biotransformation profile data in a way that was not recapitulated by 16S rRNA taxonomic structure or predicted functional pathways. Finally, expected biotransformation products and metabolic trends could be confirmed with untargeted metabolomics characterization. This scalable platform directly profiles gut microbial biotransformation activity, supporting wider applications of standardized microbiome functional phenotyping in humans and quantitative models of microbiome-competent biokinetics assessment in pharmacology and toxicology.

## Introduction

The gut microbiome alters bioavailability and bioactivities of nutrients, chemicals and drugs in the intestinal tract.^1^ While substantial variation in its composition and function affects individual responses to chemical exposures, there are a range of commonly relevant biotransformation reactions, such as hydrolysis and reduction reactions, among others. While characterization of microbial communities generally entails sequencing genetic material, measuring the metatranscriptome, proteome, and metabolome of gut intestinal contents to reveal what is present, these don’t directly quantify capacity to transform a given chemical, to what extent, or how the fermentation environment alters such activity.^2–4^ Capturing the variation and rate of biotransformation reactions across complex microbial communities is expected to contribute to better understanding gut-host effects and differences in individual responses to xenobiotic exposure.

Combining metabolomics analysis and ex vivo microbial community models has provided extensive insight into microbiota-mediated chemical metabolism. For example, Zimmermann et al. used targeted chemical screens and multi-omics approaches to map the capacity of gut bacteria to biotransform drugs, identifying numerous relevant drug-microbiome interactions.^5^ This study qualitatively linked reactions to bacteria and enzymes, but was not designed to generate biotransformation profiles to compare the functional capacities of distinct microbiota communities. In a recent report of a gut microbiota model for the analysis of bacterial modifications of xenobiotics, the authors combined the use of well-established colorimetric and dye-linked assays to profile common enzymatic activities, with HPLC and LC-MS measurements of the decay and metabolite formation from six model substrates representing key biotransformation reactions.^6^ We recently reported a chemical probe-based strategy for profiling microbiota biotransformation activity, and applied it to quantify biotransformation disruptions in the gut microbiota from rats after chemical exposures.^7^ The method provided key biotransformation metrics, but monitored only eight substrates and had limited scalability. Nonetheless, chemical probes offer a complementary strategy to profile microbiome function, including to assess multiple biotransformation activities in parallel.

In the context of reducing animal experimentation,^8^ physiologically-based kinetic (PBK) modeling is rapidly expanding as a combined *in vitro*/*in silico* method to predict biokinetics relevant to bioactivity and safety assessment.^9^ To account for gut microbial bioactivation, such models require methods to measure the kinetics of gut microbial biotransformation reactions. Batch fermentations with stool samples yield useful transformation rate data,^10^ but are limited by large bioreactor volumes and low throughput. Furthermore, growth medium,^11,12^ inoculum density,^13^ and the use of fresh versus frozen fecal material^13,14^ can alter microbial community composition and metabolic output. How these variables influence quantitative biotransformation capacity across different microbiota communities and substrates has not been investigated. Therefore, advancing ex vivo fermentation methods in predictive biokinetics requires scalable, quantitative approaches that capture microbiota biotransformation capacity and define how experimental conditions influence reaction rates.

Quantifying chemical probe depletion across time and conditions provides a functional profile of microbial biotransformation capacity. While such profiles do not replace compound-specific kinetic studies, they can reveal individual donor metabolic phenotypes, which inform how diet, drug, or xenobiotic exposures interact with microbial activity. In this study, we developed a 96-well-plate-based anaerobic fermentation workflow coupled with LC-MS/MS quantification of chemical probes to generate donor-specific biotransformation profiles. Furthermore, we assessed fermentation variables of stored fecal microbiomes for experimental reproducibility and repeated testing. The work provides a first high throughput profiling strategy to quantify biotransformations in complex human gut microbial communities and reveals how key factors in storage and growth conditions impact biotransformation rates, providing a foundation for adapting *ex vivo* fermentation to the needs of pharmacology and toxicology applications.

## Materials and Methods

### Chemicals and materials

Biotin, cyanocobalamin, p-aminobenzoic acid, folic acid, pyridoxine hydrochloride, potassium chloride, sodium chloride, ammonium sulfate (99%), magnesium sulfate, propionic acid (>99.5%), resazurin sodium salt, Amicase, yeast extract, arabinogalactan, guar gum, inulin, mucin type II, caffeine, daidzein, disopyramide phosphate, engeletin (>98%), glycochenodeoxycholic acid, isoxanthohumol, lansoprazole, lovastatin, naringin (>98%), nicardipine hydrochloride (>98%), olsalazine sodium, praziquantel, sulfinpyrazone (>99%), tacrine (>99%), terazosin hydrochloride, and sulfamethoxazole were purchased from Sigma-Aldrich. Potassium hydrogen phosphate was obtained from VWR Life Science, potassium dihydrogen phosphate and L-cysteine HCl, calcium chloride dihydrate, metronidazole, and risperidone were from VWR Chemicals. Acetic acid and sodium hydrogencarbonate were from Merck. Isovaleric acid, 4-nitrophenyl-β-D-glucuronide (>99%), hemin from bovine (98%), and isobutyric acid (>99%) were obtained from ACROS Organics. Valeric acid (>99%) was from Thermo Scientific. Soluble starch, meat extract, and arabinogalactan were obtained from Sigma-Aldrich; pectin was obtained from Alfa Aesar; xylan from Apollo Scientific. Sulfasalazine (>98%) and CUDA were obtained from Cayman Chemical, and urolithin C from abcr. LC-MS grade methanol, acetonitrile, water, and formic acid were purchased from Fisher Chemical.

### Human fecal sample collection

Consent from volunteers was collected by signature prior to sample collection. This study was exempt from review by the Ethics Committee of the Canton of Zurich. Inclusion criteria was that the donor: is in good general health, has regular bowel movements between once every 3 days to 3 times per day, has no chronic inflammatory condition of the bowel, does not use immune suppressing, blood thinning, or gut transit/ digestion-influencing medication during the month prior to sample donation, does not experience regular intestinal discomfort, and has not had an operative intestinal intervention. Anonymous fecal samples were collected directly into containers (1 L) with a sealable lid. An anaerobic packet (AnaeroGen^TM^ bag) placed into the container and sealed with the lid within 30 s after depositing the sample into the container. Donated samples were stored at 4 °C for less than 3 h before being prepared for ex vivo fermentations (**Figure 1A**).

**Figure 1.**
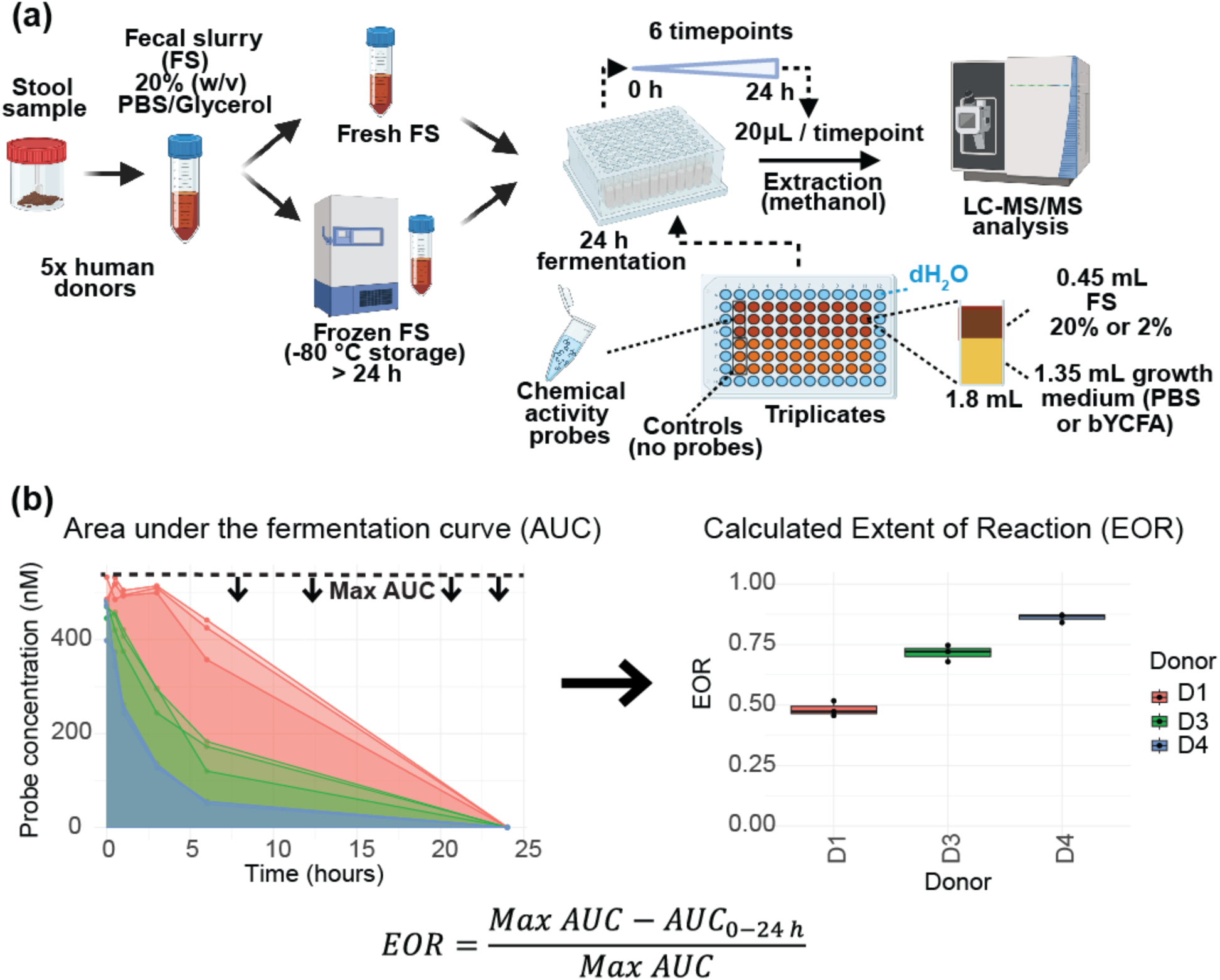
**a)** Schematic representation of post-collection fecal sample handling, fermentation assay, and chemical analysis. **b)** Two graphs summarizing the conversion of time-series concentration measurements (left plot) to EOR values (right plot). The time-concentration plot shown includes data arbitrarily selected to emphasize differential EOR values amongst donors. Each line represents one fermentation replicate. The plotted data presents data from triplicate fermentations of three donor samples under one set of growth conditions (0.5 % FS, fresh FS, nutrient media) for one chemical probe, namely glycochenodeoxycholic acid (gCDCA). The right boxplot depicts extent of reaction (EOR) values for donor-specific measurements. D = Donor. AUC = area under the curve.

### Fecal slurry preparation and ex vivo fermentations

Fecal slurry (FS) preparation and fermentations were carried out in an anaerobic chamber (BakerRuskinn Concept 500) at 37 °C with oxygen levels maintained below 0.1 % oxygen and not exceeding 2 % during equipment and sample transfer via airlock. Fecal slurries (20 % feces in PBS with 15 % glycerol) were prepared by opening the sample collection container within the anaerobic chamber, adding 10 g of stirred fecal sample to a 50 mL conical bottom tube, adding 5 5 mm glass beads, 40 mL PBS with 15 % glycerol, and shaking/vortex-mixing until the entire sample was homogenous within the slurry. To facilitate pipette transfers, the FS was passed through a coarse polyethylene sieve with 0.8-1 mm openings into a clean 50 mL conical bottom tube. The prepared slurry was then split into two 50 mL conical bottom tubes for “fresh FS” fermentation experiments or stored at −80 °C for at least 24 h before being thawed on ice for “frozen FS” fermentation experiments.

Nutrient-rich media was basal yeast casitone fatty acid media with reduced nitrogen^12^ (bYCFA) supplemented with soluble starch from potato, citrus pectin, oat spelt xylan, larch wood arabinogalactan, guar, inulin, and mucin from porcine stomach type II with oxygen removed via boiling and nitrogen gas purging. bYCFA media was made at 1.35 times concentrate to account for dilution of media with FS. Media and PBS with chemical probes were made by mixing 13.3 µL of a DMSO stock of the 20 chemical probes (**Table S1**) at 1 mM to 20 mL of PBS or 20 mL of 1.35 times concentrated rich media. Control nutrient media and PBS were made by mixing 13.3 µL of pure DMSO in PBS or media concentrate.

Ex vivo fermentations were performed in 96-deepwell plates (Abgene 96-well 2.2mL Polypropylene DeepWell plates) for 24 h. MilliQ water (2 mL) was added to the outer perimeter of the 96-well plate to reduce edge effects. bYCFA media or PBS (with or without chemical probes) (1.35 mL) was added to all fermentation wells (**Figure 1A**). Using a multichannel pipette, 0.45 mL of FS was added to each well for the given FS dilution level. Full concentration FS as described above (20 % FS:PBS w/v) was used for high inoculation conditions (5 % final FS concentration in fermentation), and diluted FS (1:9 v/v, FS / PBS with 12 % glycerol) was used for low inoculation conditions (0.5 % FS in final fermentations). Frozen fecal slurries were thawed on ice prior to fermentation. Final volume after FS addition was 1.8 mL. Immediately after FS addition, the fermentation was mixed by pipetting up and down 5 times with 0.45 mL volume. A 20 µL aliquot was immediately transferred with a multichannel pipette to a pre-prepared 96-well plate (Nunc 96-Well 450 µL Polypropylene Sample Processing and Storage Microplates with conical bottom) with 180 µL of cold extraction solvent (LC-MS grade methanol containing 400 nM sulfamethoxazole as an analytical internal standard). A gas permeable membrane (Breath Easier sealing membrane) was then adhered to the top of the plate, and the light source turned off until the next sampling timepoint. The 96-well microplate with methanol and 20 µL fermentation matrix was thermal-sealed with Easy Pierce aluminum foil, vortex-mixed 10 s, and stored at −20 °C until preparation for LC-MS/MS analysis. Samples were taken from the fermentation plate at 0 h, 0.5 h, 1 h, 3 h, 6 h, and 24 h. At each sampling timepoint the adhesive membrane was removed, the fermentation was mixed by pipetting in and out 10 times, and fermentation matrix (20 µL) was transferred to microplates with 180 µL of cold extraction solvent with internal standard as described above. The fermentation plate was then resealed with air permeable membrane and the sample plate with methanol and fermentation matrix was thermal-sealed and stored at −20 °C until further processing. After 24 h of fermentation the fermentation plates were frozen at −20 °C and material was used to make analytical controls and matrix for calibration points. The stability of chemical probes during storage was tested by preparing wells identically to fermentation experiments with either PBS or nutrient media, and chemical probes, in triplicate, then adding PBS instead of FS. Samples (20 µL) were taken as described above followed by storage at −20 °C for 15 d. Chemical probe concentrations were measured by LC-MS/MS. Stability was determined by calculating the percentage of chemical probe remaining after 15 d of storage compared to the initial concentration. Fermentations were performed in six batches over 14 d. In a first batch, we tested fresh FS prepared from donor 1 and other experimental variables (PBS/ rich media and 5 % FS / 0.5 % FS) were tested in triplicate, with and without addition of chemical probes (**Figure 1A**). In a second batch, we tested frozen FS from donor 1, and fresh FS from donor 2 and tested all experimental variables as mentioned for batch one. In a third batch, we tested frozen FS from donor 2, and fresh slurry from donor 3, and so on until the last batch was used only to test frozen FS from donor 5. In total, 240 fermentations were performed, and sampled at six timepoints resulting in 1440 samples. Calibration curve solutions were analyzed in sample matrix to account for matrix effects. Two sample matrices were made by pooling PBS or nutrient media fermentation matrix from 24 h timepoints saved from fermentation experiments. Pooled matrices were mixed with extraction solvent at a ratio of 1:9 fermentation matrix/extraction solvent (v/v) in 50 mL conical tubes, frozen at −20 °C for 24 h, centrifuged at 5000 rpm at 4 °C for 30 min, and supernatant transferred to a clean 50 mL tube. Chemical probes were diluted in these two matrices to concentrations of 1 nM, 5 nM, 50 nM, 100 nM, and 200 nM. A pooled analytical quality control (pooled QC) sample was made by mixing PBS and nutrient media matrices together at ratio of 1:1. Blanks were generated by mixing LC-MS grade water and extraction solvent at ratio of 1:9 water/ extraction solvent (v/v). Calibration curve solutions, blanks, and pooled QC solutions were aliquoted into 96-well plates and sealed with foil and stored at 4 °C in the LC autosampler prior to LC-MS/MS analysis.

### LC-MS/MS analysis

To prepare samples for LC-MS/MS analysis the sealed plates that were stored at −20 °C were centrifuged at 2850 rcf at 4 °C for 30 min. Foil seals were removed, and supernatant transferred to clean LC-MS/MS 96-well plates (SureSTART WebSeal 96-well microtiter, V-bottom, 220 µL), sealed with foil, and placed in the autosampler at 4 °C. Sample-containing 96-well plates were analyzed in a randomized order, and samples within each plate were analyzed in a randomized order. Between each 24 samples a blank, pooled quality control, and two calibration curve points were analyzed. Samples were analyzed using a Vanquish HPLC (ThermoFischer Scientific) coupled to an IDX Tribrid mass spectrometer (Thermo Scientific). A Synergi 4 µm Polar-RP 80 Å LC column (30 mm x 2 mm) RP column with SecurityGuard guard cartridge (Phenomenex) was used with a 0.7 mL/min flow rate. Mobile phases were 100 % LCMS grade water (A) and 100 % acetonitrile (B), both containing 0.1 % formic acid. The mobile phase gradient began from 0 to 0.1 minutes with 5 % B, from 0.1 to 1.8 minutes the gradient increased from 5 % to 100% B, was kept at 100 % B from 1.8 to 2.35 minutes, and then the gradient was brought from 100 % to 5 % B from 2.35 to 2.4 minutes and kept at 5 % B from 2.4-2.8 minutes followed by a 1-minute needle wash sequence prior to injection of the next sample. Each sample was analyzed in positive (+3500 V) and negative (−2500 V) heated electrospray ionization (HESI) modes as separate analyses. Other HESI source parameters are as follows, Gas Mode: Static, Sheath Gas (Arb): 60, Aux Gas (Arb): 15, Sweep Gas (Arb): 2, Ion Transfer Tube (°C): 350, Vaporizer Temp (°C): 400. Mass spectra were acquired as MS1 full scans from 60-800 *m/z*, with targeted MS/MS scans for chemical probes and internal standards (**Table S1**). MS1 scans were collected at 60k resolving power in the orbital ion trap mass analyzer with RF lens set to 50 %, maximum injection time of 50 ms, and centroided spectra. Targeted MS/MS scans were collected in the ion trap mass analyzer with optimized fragmentation in either HCD or CID (**Table S1**), quadrupole isolation window of 1.6 *m/z,* CID activation time of 10 ms, Ion trap scan rate set to Rapid, and centroided spectra. To collect nontargeted MS/MS for feature annotation, data dependent analysis (DDA) was used on a random selection of 5 % of samples throughout the sequence, and six rounds of iterative exclusion DDA^15^ were performed on pooled samples at the end of data acquisition. Two DDA MS/MS scans were collected for each MS1 scan and ions selected for fragmentation were automatically excluded for 2.5 s. DDA MS/MS scans were acquired on the orbital ion trap with mass resolution of 15k after using the quadrupole for ion isolation with window of 1.6 *m/z,* stepped HCD collision energy of 30, 45, 70, and maximum injection time of 100 ms.

### LC-MS/MS Data processing and analysis

Targeted LC-MS/MS data was analyzed with open source software Skyline^16^ version 23.1 for all chemical probes and internal standard with monitored transitions listed in **Table S1**. Summed peak areas for each targeted chemical were exported from Skyline and further data processing was performed in R version 4.6.0 (scripts available in an archived repository).^17^ To account for evaporation and sampling volume variability, the peak areas of chemical probes were scaled to antazoline which did not measurably degrade during fermentations (**Table S2**). Blank subtraction was performed by subtracting the median of triplicate fermentations without chemical probes from fermentations with chemical probes. Linear models were used to quantify chemical probe concentration and were applied to samples with the respective growth media type (PBS or rich media). The average R-squared value for all concentration curves was 0.985 (**Table S2**). Zero and missing values were replaced with levels below the limit of detection (values randomly distributed between 0.01-0.1 nM) prior to statistical analyses. One outlier measurement from one sample (chemical probe metronidazole from Donor 1 at timepoint 3 h with 5 % FS and grown in bYCFA media) was imputed to the average of the experimental triplicate due to 100x higher value than the other two fermentations (400 nM compared to 2 nM and 6 nM). Quantification error was calculated using 0 h timepoint samples with chemical probes at 500 nM (Average error was 14.3 % across all chemical probes) (**Table S2**). To quantify chemical transformation, the area under the concentration–time curve from 0 to 24 h was calculated for each fermentation. The extent of reaction (EOR) was calculated as

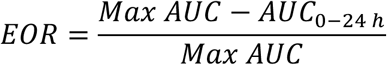

where Max AUC was defined as *C*_max_ × 24 h, with *C*_max_ representing the maximum measured probe concentration in the corresponding fermentation. EOR values range from 0 to 1, with higher values indicating greater extent of probe transformation (**Figure 1B**).

Untargeted LC-MS/MS data was processed using MS-DIAL version 4.9.221218.^18^ Data processing parameters are reported in **Table S3**. Metabolites were identified based on accurate *m/z*, and MS/MS spectral matching to a merged MS/MS spectral library of the Mass Bank of North America (massbank.us) and NIST23. Positive and negative mode ESI data were processed separately. Blank subtraction was performed by removing features with less than 10-fold maximal sample measurement divided by average of blanks. MS/MS spectral matches were manually confirmed, and in-source fragments were identified and removed when the correlation coefficient was near 1 between features at identical retention time. Adducts were combined by summation. All annotated metabolites with annotation confidence^19^ are reported (**Table S4**). Peak heights were exported for statistical analyses. Multiple comparison correction was performed using the method of Benjamini-Hochberg (BH)^20^. Clustering of metabolites was performed using Spearman correlation coefficients of metabolites across all fermentations with nutrient-rich media, with chemical probes added, followed by hierarchical clustering. The number of clusters was determined by increasing the number of clusters until metabolites with the same general trends were split into multiple clusters.

### 16S rRNA analysis

DNA was extracted (DNeasy, PowerSoil Pro Kit (Quiagen, LOT 172023704)) from the pellets that remained after supernatant was removed from the microtiter 96-well plates. The contents of wells corresponding to timepoint zero in PBS growth medium were pooled for DNA extraction for each donor. The V4 region of the 16S rRNA gene was amplified using 515F/806R-region primers and sequenced using paired-end 300-bp Illumina sequencing. Demultiplexed raw sequences were processed using QIIME 2 (v 2024.10)^21^. Reads were imported into QIIME 2 and denoised using the DADA2 plugin, which included quality filtering, dereplication, chimera removal, and paired-end merging. During denoising, sequences were truncated at 260 bp (forward) and 200 bp (reverse) based on quality score profiles and reads with ambiguous bases were removed. Merged reads were required to be at least 260 bp in length. Amplicon sequence variants (ASVs) were retained only if they passed chimera filtering and were present in at least one sample. Taxonomy was assigned with a Naive Bayes classifier trained on the Greengenes2 2024.09 backbone V4 reference at 99 % similarity, specific to the 515F/806R region. ASVs for family (L5) and genus (L6) levels were exported for downstream taxonomic analysis. Functional predictions were performed using PICRUSt2 (v 2.5.2)^22^. Gene family abundances (KEGG Orthologs, EC numbers) were predicted, and MetaCyc pathway abundances were inferred from predicted gene family content. Downstream integration with chemical data was conducted in R^17^. Taxonomic groups, functional profiles (KO, EC, and MetaCyc pathway abundances) were converted to relative abundances and removed if the relative abundance was below 0.0001. Chemical EOR data were averaged across replicates for each donor and compared with taxonomic groups or microbial functional profiles using Spearman correlation.

## Results

### Variations in extent of reaction of chemical probes depending on structure

Microbial metabolic activity was profiled by measuring the extent of transformation of 20 chemical probes. The panel included pharmaceuticals, diet-associated compounds, and endogenous metabolites selected to report established or anticipated gut microbial reactions including deglucuronidation, deglycosylation, bile acid deconjugation, azo-bond reduction, sulfoxide reduction, nitroreduction, and benzisoxazole ring cleavage, together with compounds lacking reported microbial transformations as a reference for minimal expected microbial activity (**Figure S1 A**). To capture diverse potential profiles of reaction progression, we calculated a normalized reaction rate metric (hereafter referred to extent of reaction (EOR)) using the area under the curve (AUC) of concentration plotted over 24 h for each individual fermentation resulting in EOR metric from 0 to 1 with 1 being the highest, and 0 being lowest EOR (**Figure 1B**). Here, EOR encompasses all chemical transformations the probe undergoes, both enzyme-catalyzed and spontaneous. Prior to settling on EOR as a useful metric, we modeled degradation kinetics assuming exponential decay and calculated first-order rate constants (k). While the degradation profiles of several probes fit this model well (R^2^ > 0.9), others had low R^2^ measures and highly variable fits, consistent with higher-order kinetics (**Figure S1 B**). This variation likely arises from probe-specific biochemistry, matrix interference, and/ or non-saturated enzyme conditions.^23^ Exponential decay rate constant k and AUC-based EOR both reflect reaction velocity and are visually compared showing reaction profiles following first order kinetics (**Figure S1 C**). To determine the extent of spontaneous probe degradation, the percentage of remaining probe after 15 d storage at −20 °C in extraction solvent, plus PBS or bYCFA matrix was measured (**Table S2**). For most probes, >90% of the original concentration remained after 15 d; exceptions were disopyramide (85%), isoxanthohumol (81%), lansoprazole (87%), lovastatin (73%), and praziquantel (79%), which had the indicated remaining fraction of concentration (**Table S2**). EOR varied greatly depending on probe structure. For some probes (e.g. nitrophenyl-glucuronide and gCDCA), most EOR values exceeded 0.9. For other probes, (e.g. disopyramide and praziquantel) most EOR values were below 0.15 (**Figure 2**). In this manner it was possible to compare fermentations using EOR for each individual probe, and overall EOR profiles.

**Figure 2.**
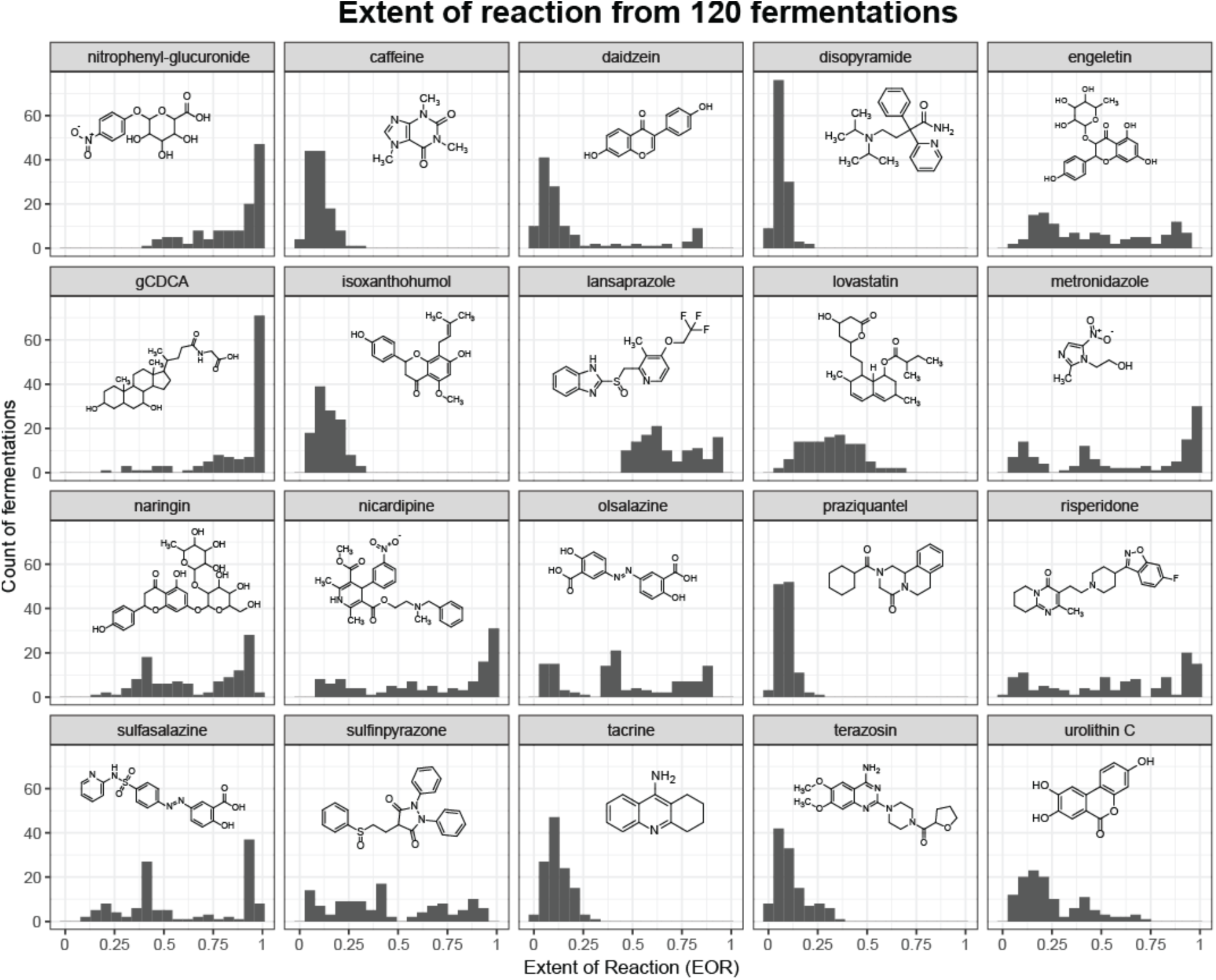
Histograms of EOR values for each chemical probe and their structures. Data are from triplicate fermentations of all treatment conditions for all donors (n=120). Bin width of the histogram is 0.05 on x-axis. gCDCA = glycochenodeoxycholic acid.

### Fermentation conditions influence biotransformation profiles and allow discrimination amongst donors

Having established a robust method to profile variations in biotransformation activities in microbial communities, we aimed to determine the impact of common condition variables relevant to ex vivo fermentation modeling, namely inoculation level, media type, and whether samples were processed fresh or frozen. For this evaluation we determined EOR as described above and evaluated differences using mixed linear effect models (LMMs) with microbiota donor as a random variable and inoculum level (5% or 0.5% FS), media type (nutrient media or PBS), and storage (fresh or frozen FS) as fixed variables. Changing FS inoculum concentration altered the EOR for 17 of the 20 chemical probes. Of these, 16 EORs increased at the higher FS inoculum (5%), yet decreased for lovastatin (**Figure 3A, Table S5**). This general increase was expected due to the conditions favoring microbial growth and thus the presence of higher enzyme levels and activity. Lovastatin may be an outlier because it spontaneously degrades more than any other probe (**Table S2**), consistent with its known propensity to degrade in water, particularly under basic conditions.^24^ In PBS vs nutrient media, 9 probes degraded faster in nutrient media, consistent with nutrient availability promoting microbial load and enzymatic activity. Nonetheless, nitrophenyl-glucuronide, urolithin C, and terazosin degraded faster in PBS (**Figure 3B, Table S5**), potentially due to altered metabolic regulation in response to nutrient scarcity^25,26^, or competitive inhibition by nutrient media components not present in PBS. Finally, despite the expectation that microbiome freezing alters microbiota viability and composition after re-inoculation^14,27^, comparing fresh vs. frozen FS did not suggest statistically significant alterations, and biotransformation capacities are generally stable after freezing with 15% glycerol (**Figure 3C, Table S5**). Consequently, this observation justifies the storage of fecal slurries at −80 °C as a reliable preservation method for future large scale microbiome biotransformation profiling.

**Figure 3.**
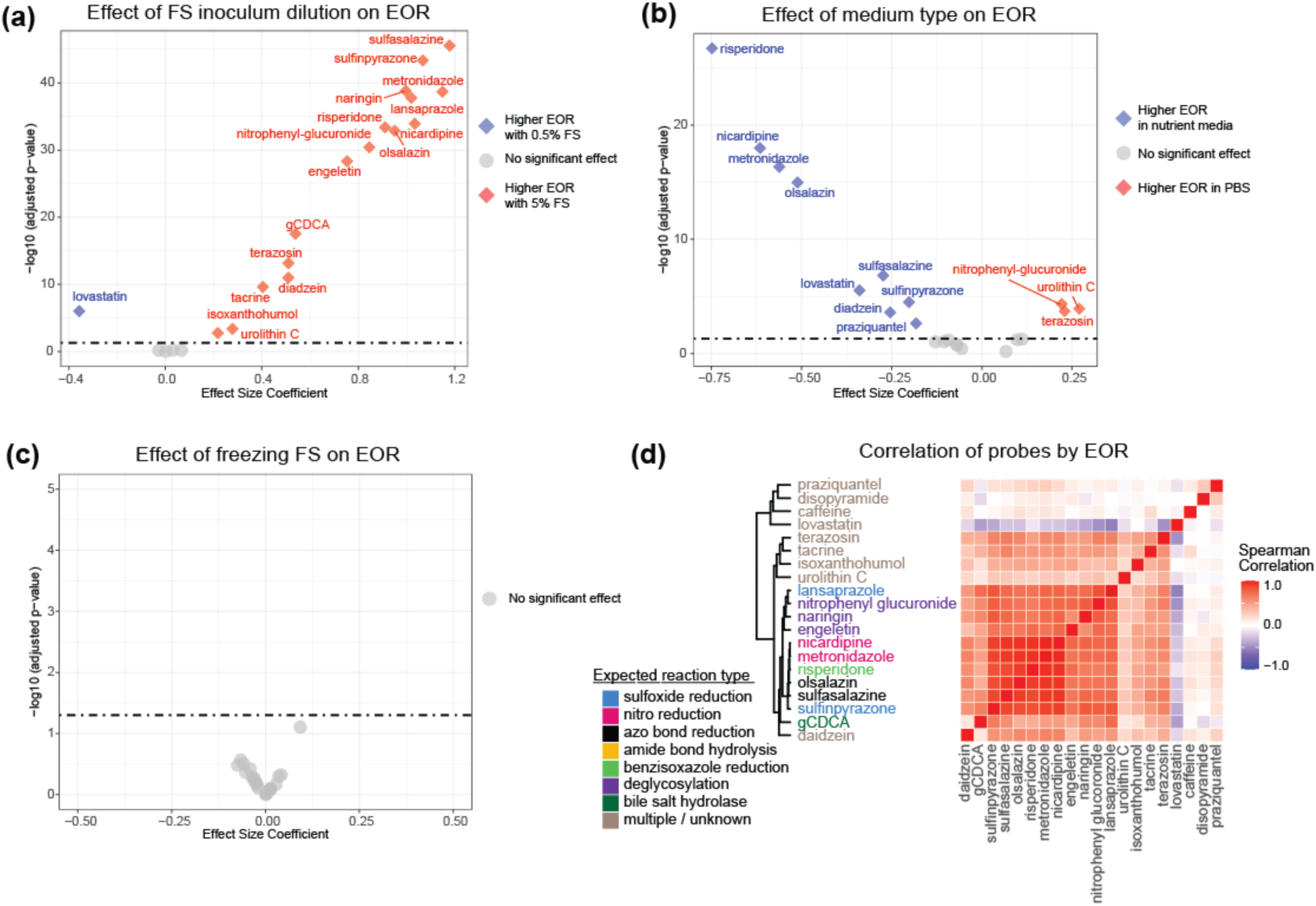
Effect of fermentation conditions on EOR for all probes on the basis of linear mixed effect models. **a)** FS dilution level (5% vs. 0.5% FS inoculum). **b)** Growth medium (nutrient media bYCFA vs. PBS) **c)** Fresh compared to frozen FS. Benjamini-Hochberg method was used to adjust p-values for multiple comparisons. **d)** Correlation of EOR and probe identity for probes in all fermentations. The expected chemical reaction type for each probe is linked by color.

Consistent with close hierarchical clustering of correlation coefficients for all chemical probes (**Figure 3D**), we anticipated that probes which undergo similar reactions (**Figure S1A**) would have similar EOR trends after perturbations. Nonetheless, low EOR values (less than 0.5) also generally clustered together on the top half of the correlation matrix (**Figure 3D**), and similar correlation was observed also for different chemical reaction types, so microbial activity in general may be driving the correlations between probes. This case of multicollinearity also complicates association of EOR values to non-targeted metabolite features or predicted microbial function values from genomic analyses.

Having established the individual probe trends, we next assessed how samples from five different donors may be differentiated based on biotransformation profiles by PCA of EOR measurements. To account for the large effect of fermentation conditions, samples were separated by inoculation level (5% and 0.5%) and media type (PBS and nutrient rich) resulting in four condition-specific groups (**Figure S2**). We observed donor-specific clustering with high FS inoculation (5%), regardless of media type **(Figure S2A, S2B)**. Donor 4 was the most strongly separated donor, and had consistently lower EOR values for risperidone, olsalazine, sulfinpyrazone, and metronidazole **(Figure S2A, S2B)**. In contrast, PCA from low inoculation (0.5%) conditions resulted in reduced separation between donors, consistent with overall lower EOR values (**Figure S2C, S2D**), but there were donor-associated trends for gCDCA and 4-nitrophenyl glucuronide that were not evident with 5% FS inoculation. For example, the EOR of gCDCA is near 1 in all 5% FS fermentations (either PBS or nutrient media, **Figure S2 A and B**), while adjusting the FS to 0.5% lowered the gCDCA EOR for donors 1 and 3 more than for other donors (**Figure S2 C and D**). We therefore built a hybrid EOR profile by selecting, for each probe, the fermentation condition with the greatest EOR variation across samples (**Figure 4A; Table S2**). This reduced the effect of saturating the upper limit of EOR values and improved donor separation by PCA (Figure 4B), indicating that donor-level biotransformation profiles are best resolved when each probe is measured under conditions that place it within an informative dynamic range.

**Figure 4.**
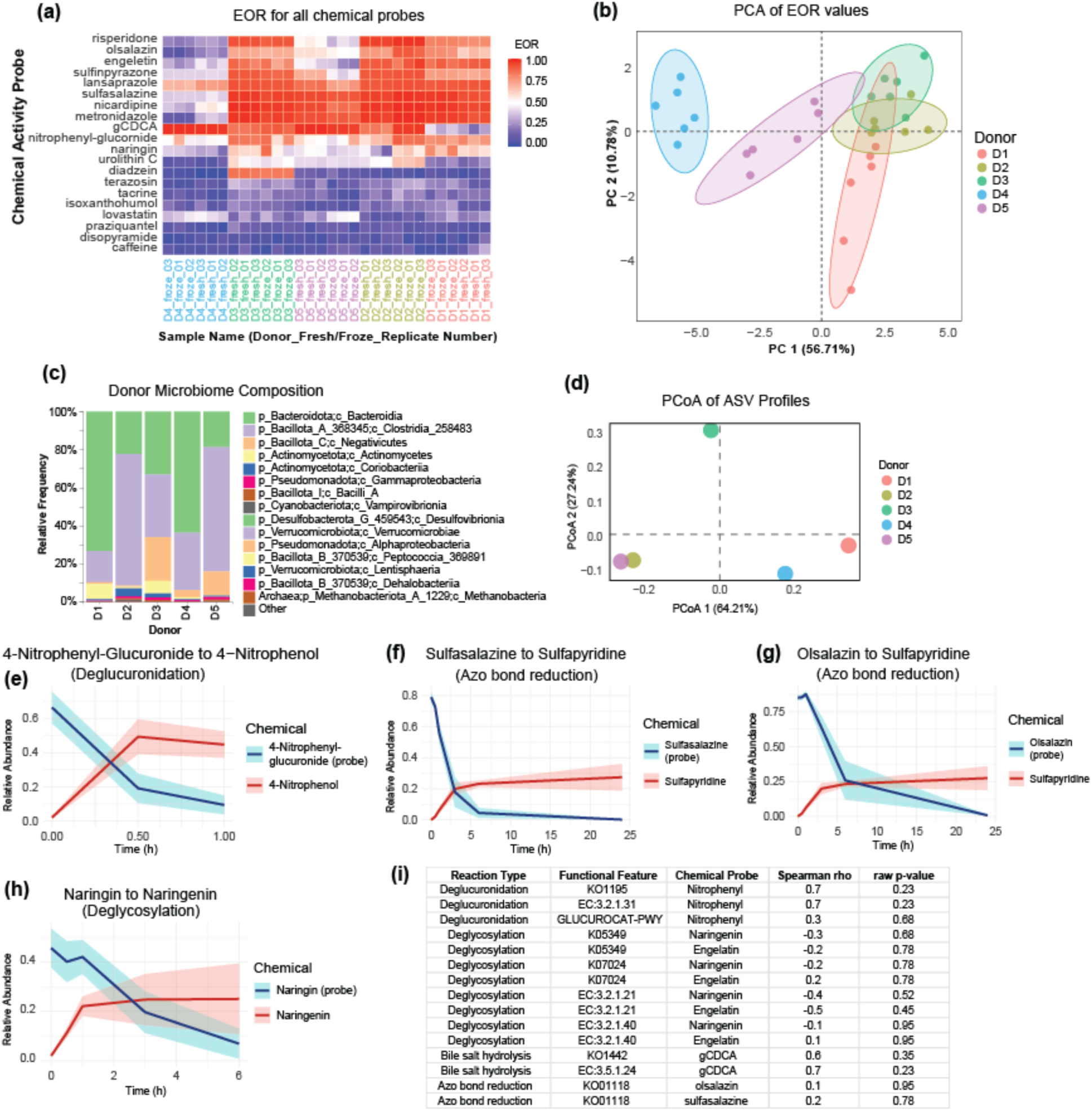
**A.** Heatmap of EOR values for each chemical probe for all samples. Data is displayed for conditions that maximized variation across samples (**Table S2**). Both x and y axis are arranged by hierarchical clustering (hclust; median agglomeration). **B.** PCA score plot from EOR values and each sample colored by donor number. Shaded ellipses represent 90 % confidence interval around each donor. **C.** Taxonomical stacked bar chart of relative abundance of class level (L3) for each donor sample. **D.** PCoA from relative taxonomic abundances with color representing donor number. **E-H.** Ribbon charts of chemical probe (blue) and expected metabolite (red) relative abundance over time from fermentations in nutrient media with 5 % FS inoculation. **I.** Table of correlation test results (Spearman, n=5) between inferred functional features from Picrust2 to EOR values for the indicated chemical probe (mean value from six replicates from panel A). Functional features were manually curated to match expected reactions from chemical probes and designation of KO indicates the feature is from KEGG, EC indicates it is a feature from the Enzyme Commission, and -PWY indicates the feature is a MetaCyc pathway. No correction for multiple comparisons was applied to these *p*-values.

To determine the most sensitive probes for differentiating donor samples based on biotransformation profiles, partial least squares discriminate analysis (PLS-DA) was used followed by variable importance in projection (VIP) ranking (**Figure S3 A,B**). The PLS-DA model had moderate classification performance (R^2^Y = 0.47, Q^2^Y = 0.42, p = 0.05) and, as expected, separated donors more fully than PCA, supporting its use to identify the most discriminating probes for donor differentiation. The top 4 most discriminating probes were gCDCA, urolithin C, daidzein, and engeletin with VIP scores above 1 (**Figure S3 B**). Both bile acid metabolism^28^ and daidzein biotransformation^29^ are known to vary greatly amongst individuals making it logical that these probes are so effective. The chemical probes with the lowest VIP scores (below 0.75) were disopyramide, caffeine, and isoxanthohumol, which contribute minimally to differentiating donors. These three chemicals had some of the lowest average EOR of all probes, indicating that the conditions used in these fermentations were not ideal for these transformations to occur. In the future, adjusting fermentation conditions may increase the rate of biotransformation for these probes and allow valuable metabolic insight.

### Biotransformation profiles offer a unique basis of characterization compared to taxonomic composition

Having established the performance of biotransformation profiles to differentiate donors, we then addressed how community profile analysis relates with EOR values. We used 16S rRNA profiling to analyze the microbiome of the 5 donor samples (**Figure 4 C**) and performed PCoA to assess whether groupings derived from overall community structure mimicked relationships derived from EOR values. PCoA distribution indicated that the community of donor 3 was the most unique sample, while donors 2 and 5 were highly similar (**Figure 4 D**), whereas EOR based PCA plots indicated that donor 4 was the most unique, while donor 3 was similar to donors 1 and 2 (**Figure 4 B**). We then investigated how predicted functional activity for probes with known metabolism pathways in KEGG, EC, and/or MetaCyc databases related with EOR values. These included deglycosylation of 4-nitrophenol-glucuronide, naringin, and engeletin, hydrolysis of glycine from gCDCA, or azo bond reduction of sulfasalazine and olsalazine (**Figure 4 E-H**). To confirm the anticipated reactions, metabolites were identified in the non-targeted metabolomics data (**Figure 4 E-H**). Apparent increases in metabolite abundances support their relevance as the basis of probe transformation, but they were not quantified so other competing potential reactions could not be assessed. There were no statistically significant correlations between predicted functional activity and experimentally measured EOR values for any of the 5 donor samples for any of the 13 feature-probe relations tested (**Figure 4 I**). This is in agreement with a previous observation that metabolic gene presence does not strongly correlate to corresponding metabolites in stool or blood serum even with a sample size of 1,135 human donors^3^ and demonstrates that biotransformation profiling captures information that cannot be accurately inferred from community composition or predicted pathway abundance.

### Metabolome-wide variations are individual and dynamic

Given the large differences in microbial community characteristics between taxonomic composition and biotransformation profiles, we further investigated metabolism trends metabolome wide. For this purpose, non-targeted LC-MS/MS analysis was used to detect 7855 chromatographic features above the level of blanks. After manual curation, 337 level 2 (accurate *m/*z plus MS/MS library match) or higher^19^ unique metabolite annotations were retained (**Table S4**). Pooled QC replicates clustered tightly by PCA (**Figure S4 A**) and the median percent relative standard deviation (% RSD) of metabolites in pooled QC replicates was 11.9 %. Growth medium type and FS conditions were the primary determinants of clustering in PCA score plot (**Figure S4 A**), and sampling timepoint systematically influenced PCA clustering (**Figure 5 A-D**). Nutrient rich media conditions resulted in temporal metabolome shifts (**Figure 5 A,B**), while PBS was the basis of little or no clustering of timepoints (**Figure 5 C,D**). FS inoculation of 5 % or 0.5 % influenced how quickly the metabolome shifted in nutrient-rich media (**Figure 5 A,B**). Inoculation rate also revealed donor-specific metabolome profiles that can be observed by separate clustering of donor 4 by PCA in the 5% FS inoculation groups (**Figure 5 A,C**), but little or no interindividual clustering in 0.5% FS inoculation groups (**Figure 5 B,D**). These trends can be explained largely by the high abundance of energy-containing metabolites in nutrient media that are transformed by microbial fermentation over time, thus shifting the metabolome, whereas PBS is nutrient scarce and does not induce a metabolome shift over time. The more rapid metabolome shift in 5% FS inoculum compared to 0.5 % FS inoculum in nutrient media is expected to be due to larger microbe content causing faster nutrient metabolism compared to 0.5% FS.

**Figure 5.**
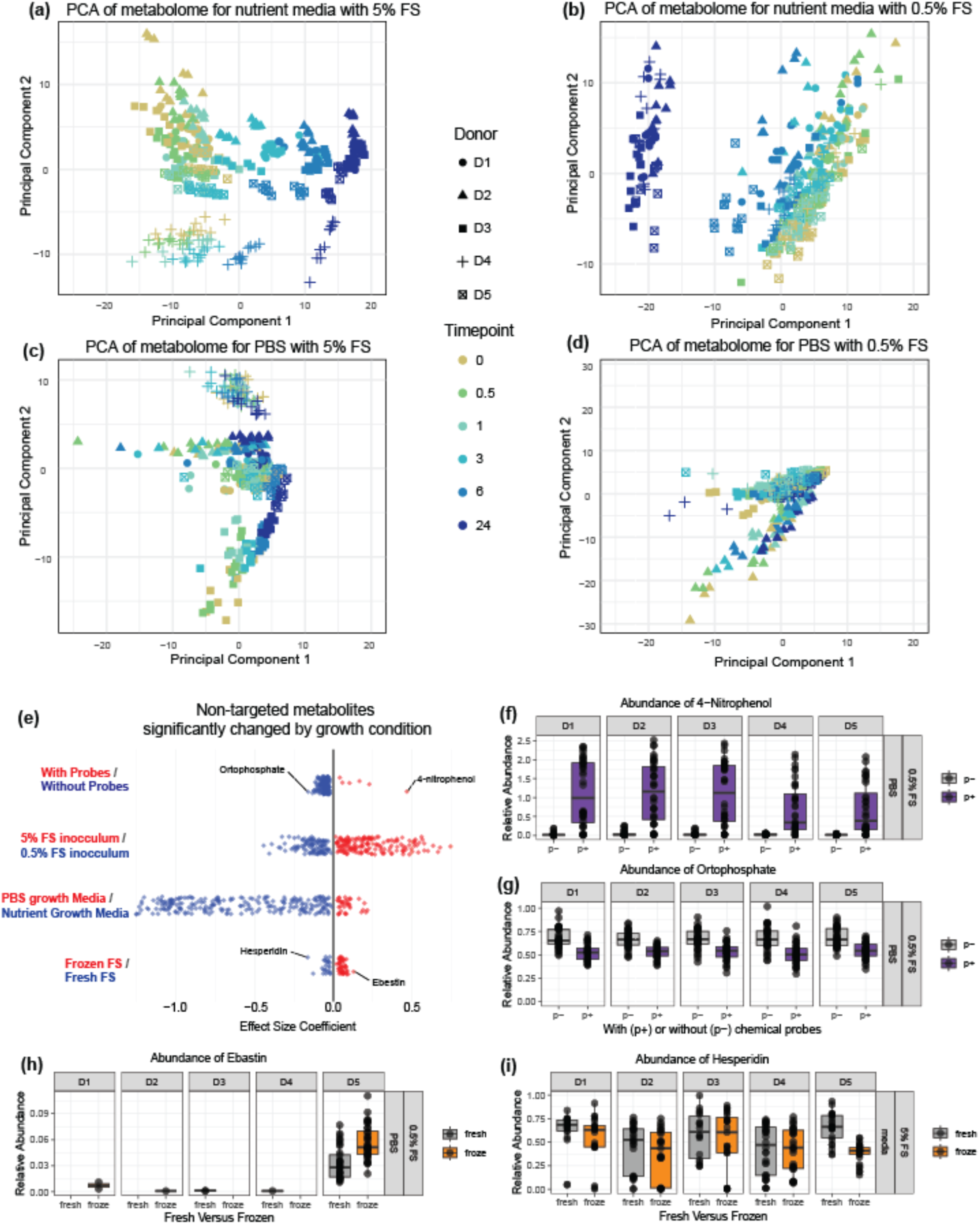
PCA score plots of 24 h fecal fermentations amongst 5 donors. Data are filtered non-targeted metabolite levels (n=312 metabolites) from samples grown in **A**. nutrient media with 5 % FS inoculation, **B**. nutrient media with 0.5 % FS inoculation, **C**. PBS with 5 % FS inoculation and **D**. PBS with 0.5 % FS inoculation. **E**. Metabolites significantly affected by growth conditions based on linear mixed effect model with growth conditions and timepoints set as fixed variables and donors as random variable. Each point represents a metabolite significantly (p < 0.05; BH adjusted n=1812 tests) affected by the growth condition indicated. Growth condition label color (red/ blue) indicates which metabolite points are increased in that condition. Four metabolites are labeled and further plotted as boxplots to investigate the effect of chemical probes on fermentation **F-G**, or affected by freezing **H-I**. Boxplots display a subset of data indicated by panels on the right side of each plot with all data plotted in associated supplemental figures S5.

To identify metabolites that respond to the tested fermentation conditions, linear mixed effect models were generated for each of the non-targeted metabolites. In this case, an additional fixed variable of whether probes were added to the fermentation was tested. When probes were present during fermentation, 170 metabolites decreased and 6 metabolites increased compared to control fermentations without probes present (**Figure 5E, Table S7**). The largest effect size coefficient under these conditions was for 4-nitrophenol, explained by formation of 4-nitrophenol from 4-nitrophenol glucuronide (**Figure 5F**). Metabolites with decreased abundance when chemical probes were added resulted in relatively lower effect size coefficients with the largest effect appearing in orthophosphate (**Figure 5G**). To determine the extent of the effect of probes on fermentation metabolites, PCA was used to visualize samples grown in nutrient media with 5 % FS inoculum and revealed no clustering by probe presence within each timepoint cluster (**Figure S4 B**). Therefore, we conclude that chemical probe addition has a small enough effect on metabolism to allow EOR profiles to represent biologically relevant metabolism phenotypes of microbiota.

Comparing FS inoculum levels, 5 % FS corresponded to an increase in 164 metabolites and decrease of 103 metabolites compared to 0.5 % FS (**Figure 5E**). The effect size coefficient was much larger for metabolites that were higher in the 5 % FS conditions, and the most strongly affected metabolites included bile acids and diet-related metabolites expected to be derived from the FS itself. All 103 metabolites that were higher in the 0.5 % FS were also significantly higher in nutrient media compared to PBS, leading us to conclude that these media components are metabolized by the microbiota.

For nutrient media comparison to PBS, there were 227 metabolites higher in nutrient media and 37 metabolites significantly higher in PBS (**Figure 5E**). The complex nature of nutrient media explains the large proportion of metabolites that are higher compared to PBS. Of the 37 metabolites detected at higher levels in PBS compared to nutrient media, 34 of them are higher in 5 % FS inoculum compared to 0.5 % FS inoculum indicating they are FS derived (**Table S7**). One explanation of this observation is more ion suppression during LC-MS analysis of ion rich nutrient media samples compared to PBS samples resulting in higher abundance values in PBS containing fermentations.

For fresh vs frozen FS fermentations, 24 metabolites were more abundant in fresh FS, and 49 metabolites were higher in frozen FS. More than 90 % of these metabolites were also higher in 5 % FS inoculation compared to 0.5 % FS inoculation indicating they are FS-derived metabolites (**Table S7**). The metabolite with the highest increase in frozen samples compared to fresh FS fermentations was ebastine (**Figure 5H**), an antihistamine drug. One explanation for the increased level of metabolites in frozen fecal samples is release of chemicals bound to insoluble material during a freeze-thaw cycle. Hesperidin and neohesperidin, two glycoside containing plant metabolites from the nutrient media, had the largest effect size increase in fresh FS (**Figure 5I, Table S7**), possibly due to more glycosylase activity towards these substrates in frozen samples. Of note, the effect size coefficient here was relatively small and driven by a single donor (**Figure 5 H, I**), therefore limiting possible generality.

To investigate global metabolite dynamics during fermentation we focused on the conditions with the strongest time-dependent shifts, namely 5% FS inoculation in nutrient-rich media (**Figure 5A-D, Figure 6A**). To group metabolites by similar temporal abundance patterns, we used hierarchical clustering of Spearman correlations. The number of metabolite clusters was chosen using the gap statistic that supported the presence of six clusters and was further supported based on biological interpretability and diminishing gains in gap score.^30^ Manual inspection revealed clear temporal patterns within clusters 1, 3, and 6, whereas clusters 2, 4, and 5 were driven more by interindividual variation (**Figure S6A**). Cluster 1 metabolites generally decreased over time and likely represent media or FS-derived substrates consumed during fermentation, including sucrose and the dipeptide Ile-Leu (**Figure 6B**). Cluster 3 metabolites increased over time and included known microbial fermentation products such as lactic acid, 3-phenyllactic acid, hydroxyisocaproic acid, hydroxybutyric acid, pantetheine, and 4-vinylphenol, supporting their assignment as microbially generated metabolites (**Figure 6C**).

**Figure 6.**
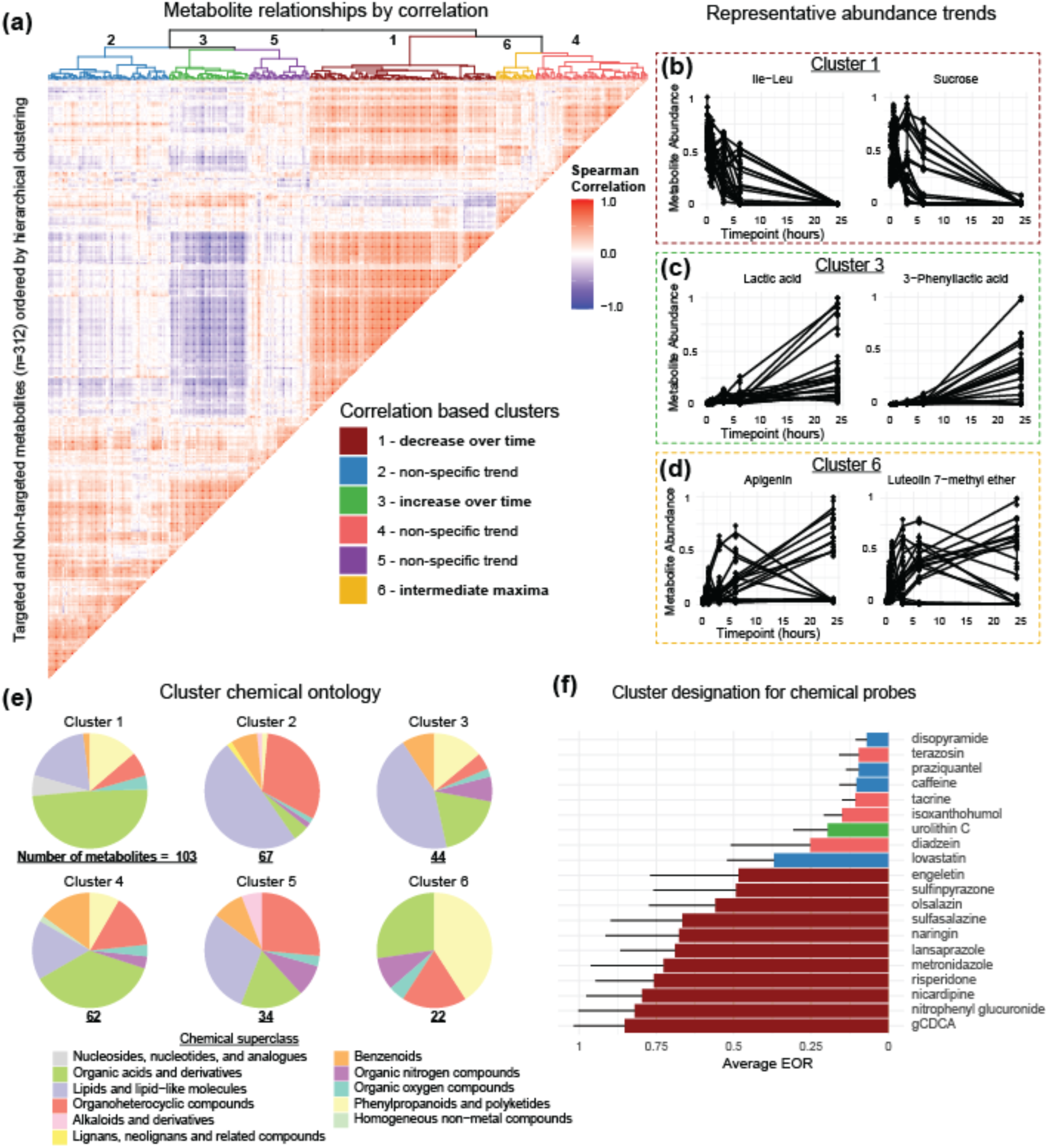
**A.** Spearman correlation matrix of metabolite abundances in fermentations performed in nutrient rich media, and with chemical probes added. Metabolites with more than 20 % missing values were removed from analysis. Hierarchical clustering was performed using agglomeration method Ward D2 to order metabolites and perform clustering. **B-D.** Two representative metabolite trend plots from three clusters of interest are displayed. Metabolite abundance is normalized peak heights scaled between 0 and 1. **E.** Chemical superclass ontology calculated by ClassyFire displayed as pie charts for each metabolite cluster including number of metabolites below each cluster plot. **F.** Bar chart indicating the average EOR of chemical probes from fermentations performed in media, with color indicating which cluster the probe is located (see legend from panel A). Order of probes is determined from lowest to highest average EOR.

Cluster 6 metabolites increased over time in some samples and increased then decreased over time in others. Metabolites in this cluster include flavonoid intermediates such as apigenin and luteolin 7-methyl ether, as well as succinic acid and choline (**Figure 6D, Figure S6B**). In 5% FS fermentations, these metabolites peaked early (0.5-3 h) and then decreased, whereas in 0.5% FS fermentations they continued accumulating over 24 h, consistent with intermediate products in metabolic pathways that are further transformed as fermentation proceeds.

These clusters were also characterized using ClassyFire^31^ based chemical ontologies (**Figure 6E**). Cluster 1 had the highest proportion of organic acids and peptides compared to all other clusters, including 42 unique di- and tri-peptides and all six nucleoside/nucleotide metabolites, indicating rapid microbial uptake of these growth supporting substrates for protein or DNA synthesis^32^. Cluster 2 contained the highest proportion of lipid and lipid-like molecules that includes 10 detected bile acids, and also the only lignan enterolactone. These metabolites are microbially transformed indicating that cluster 2 contains a portion of microbially derived metabolites even if they do not follow the increasing trend of cluster 3 (**Figure 6 C**). Cluster 3 also contains a large proportion of lipids and lipid-like molecules, including relatively small (<200 Da) microbially produced lipids 2-Hydroxy-2-methylbutyric acid^33^ and azelaic acid^34^, in addition to 7 glycerophospholipids, a major component of bacterial lipid membranes^35^. Cluster 6 contained the highest proportion of phenylpropanoids and polyketides, consistent with its enrichment for flavonoid intermediates and was the only cluster devoid of lipids. Overall, chemical ontology supports the interpretation that temporal clusters reflect different areas of metabolism such as substrate utilization, product formation and intermediate turnover.

### Linking chemical probe EOR values to non-targeted metabolites

Given apparent dynamic trends revealed by clustering analysis of metabolome-wide data, we evaluated the corresponding behavior of the chemical probes. Thus, eleven of the chemical probes fall into cluster 1 in agreement with the trend of this cluster decreasing over time. The other 9 probes were split between clusters 2, 3, and 4. We tested whether probe EOR values played a role in determining its cluster and found that all 11 probes in cluster 1 had higher average EOR than probes in other clusters. Urolithin C unexpectedly lands in cluster 3 where metabolites have a general increasing trend, but after further investigation we found that 2 donors have slight increases in urolithin C concentration. Urolithin C is a known microbial product of ellagic acid metabolism, a chemical found in a variety of fruits, indicating that some donors likely ate food containing ellagic acid, and during ex vivo fermentation this ellagic acid was converted to urolithin C causing it to increase. We also examined expected products of chemical probe transformations in the untargeted LC-MS/MS data beyond the reactions indicated previously (**Figure 4E-H**). Several putative annotations increased over fermentation time and were enriched in probe-containing samples, supporting the occurrence of the expected microbial transformations across the probe panel (**Figure S1A, Figure S7**).

We next investigated how chemical probe EOR values and non-targeted metabolite levels correlated in stool samples. Setting stool baselines corresponding with PBS growth conditions and 5 % FS inoculum at timepoint zero, 115 metabolites significantly correlate to at least one chemical probe (p<0.001, BH correction for n=6820 comparisons) (**Figure S8 A**). About one third of stool metabolites correlate to the EOR values of just one chemical probe, while another third of metabolites are correlated to more than 6 chemical probes (**Figure S8 B**). For example, the fecal metabolite stercobilin is significantly positively correlated with 12 chemical probes, and negatively correlated with lovastatin (**Figure S8 A, C**). Stercobilin is a microbially generated tetrapyrrole with a specialized enzyme expected to be necessary for formation^36^, and the 12 probes are not chemically similar or expected to be insightful into the enzymatic formation of stercobilin, but rather indicative of total microbial metabolism occurring in the sample. We observed high degrees of internal correlation within both metabolite profiles (**Figure 6A**) and EOR measurements (**Figure 3D**), indicative of multicollinearity. Additionally, three chemical probes (praziquantel, disopyramide, and caffeine) had no significant correlations to a non-targeted metabolite. These three chemical probes also have the three lowest average EOR values across all fermentations, suggesting minimal transformation over 24 h, and therefore no biological association with chemicals within the original stool samples. Therefore, correlations between metabolites and probes were not sufficient to link chemical probes to specific non-targeted metabolites, but relevant relationships may emerge in future evaluation of larger sample size and diversity.

## Discussion

We report here the use of a panel of chemical probes to profile intestinal microbe-catalyzed reactions as a basis of stratifying individual donor samples by biotransformation activity, and to understand how this functionality is influenced by ex vivo fermentation conditions. Across 20 probes, extent of reaction (EOR) varied strongly by probe structure and fermentation condition, demonstrating that biotransformation capacity is both chemically specific and dependent on the fermentation environment. Inoculum level and growth medium had the largest effects on EOR profiles, whereas frozen fecal slurries preserved biotransformation activity relative to fresh samples, supporting their use for repeated ex vivo profiling. When probe-specific conditions were selected to maximize informative variation, EOR profiles separated donor microbiomes by PCA and revealed functional differences not captured by 16S rRNA community structure or predicted pathway abundance. Untargeted metabolomics further showed that fermentation produces coordinated, donor- and condition-dependent metabolome shifts. Together, these findings establish biotransformation profiling as a direct, scalable readout of microbiome functional capacity that complements sequence and metabolome focused approaches.

Given the highly dynamic nature of microbial biotransformation kinetics, and dependence on ex vivo fermentation conditions, there is no single/standard method to capture vast reaction rate ranges. Nonetheless, the result of this study suggests experimental design best practices for quantitative biotransformation assessment. For example, as expected, diluting FS inoculum generally reduced EOR for reactions like nitroreductions and deglucuronidation carried out by microbial enzymes, highlighting that controlling inoculum density is crucial. Thus, quantitative measurements should be performed within the linear range of the dilution-rate relationship where substrate conversion rates are proportional to biomass. With regards to the influence of media composition on biotransformation, we found that nutrient-rich medium versus a minimal buffer affected metabolic activities in different ways. For example, using a nutrient rich growth medium (bYCFA+6C+MUC) significantly increased EOR values for putative transformations like benzisoxazole ring reduction, nitroreduction, and sulfoxide reduction relative to nutrient-scarce buffer (PBS), whereas other activities like deglucuronidation and primary bile acid deconjugation were the opposite, suggesting enzyme-specific differences in sensitivity to activity induction with nutrient availability. Thus, medium-induced effects are chemical probe-dependent, and researchers must carefully account for these nutrient-driven effects when designing biotransformation assays. Finally, using frozen fecal samples that were stored at - 80°C in 15 % glycerol, yielded metabolic profiles and EOR values that were statistically indistinguishable from fresh samples over the 24-h fermentation period examined. This aligns with evidence that appropriate cryopreservation can preserve microbiota functionality for ex vivo studies.^14,27,37^ The storage comparison tested one cryoprotectant condition and short-term freezing, therefore it would be useful to conduct future experiments to assess longer-term biobanking and repeated freeze-thaw effects. Nonetheless, these results are highly encouraging with regards to the reproducible use of stored fecal slurries for repeated biotransformation testing with the same microbiome, which is important for biokinetic studies in pharmacology and toxicology.

Stratifying individual donors based on their biotransformation phenotypes provides a foundation for resolving interindividual variability and uncovering distinct functional cohorts but depends on profiling donors under suitable conditions. High inoculum conditions generally improved donor separation, whereas lower inoculum conditions revealed additional donor-specific differences for reactions that saturated under high-biomass conditions. By creating a hybrid EOR profile, i.e. selecting probe data best stratified by conditions, donors better separated by PCA. These results support the capacity to analyze microbiomes as chemically active systems, not only as taxonomic communities. The donor set was small here, suggesting in future studies the analysis of larger cohorts to address population-level variability, reproducibility, and exposure-relevant effect sizes. Indeed, in this work, there was a weak association between 16S rRNA-based community structure, predicted functional activity, and EOR profiles. Although expected products were observed for several probe reactions, we could not identify predicted functional features that correlated significantly with EOR values for five donors. These observations highlight the inequivalence of gene presence, predicted pathway abundance, and chemical transformation capacity, and the current limitation in effectively linking these under defined assay conditions.^3,4^

In addition to the targeted measurements, the comparative untargeted metabolomics analysis performed with the same samples provided a global view of microbiota-driven chemistry during fermentation. We detected and putatively annotated hundreds of features, revealing metabolic interactions that depended on both growth conditions and the donor’s microbiome.

Untargeted metabolomics data was also valuable to confirm the occurrence of several expected chemical probe biotransformations. However, due to the qualitative nature of untargeted LC-MS analysis, the concentration of products was not directly measured, making it impossible to dissect contributions of other competing chemical reactions. Indeed, a limitation of using EOR is that the metric integrates probe loss due to microbial biotransformation, as well as spontaneous degradation, matrix effects, and unmeasured off-target chemistry. This is especially relevant for probes that can spontaneously degrade, which could be addressed in the future by quantifying enzymatically produced probe products. The time-series sampling strategy enabled correlation analysis to provide insight into chemical clusters that could identify microbially generated metabolites, offering an additional advantage as this information cannot be found when only analyzing stool samples. Correlations between baseline stool metabolites and probe EOR values were similarly informative but biologically inconclusive because many metabolites and probes were internally correlated, limiting assignment of individual metabolites to specific reaction mechanisms. Larger studies with broader donor diversity may make these relationships more meaningful.

Overall, this ex vivo microbiota biotransformation profiling platform represents a novel and practical tool for quantitative measurements of multiple biotransformations simultaneously, which is a key advance for translating microbiome data into pharmacokinetic and toxicological predictions. Compared with cultivation methods focused primarily on community composition and general metabolic outputs like short chain fatty acids,^12^ this approach specifically quantifies transformation rates of defined chemical probes and maps them onto known and putative metabolites. While the probe panel covers multiple reaction classes of high relevance to gut microbial reactions, it is by no means exhaustive, and probes with slow reaction progression such as caffeine, disopyramide, and praziquantel may require adjustment of concentration, incubation time, inoculum densities, or analytical approaches to provide meaningful data. Additionally, fixed time batch fermentation format is a simplification of the intestinal environment but does not account for host absorption, spatial structure, transit time variation, immune signaling, or continuous substrate supply. Future studies incorporating absolute product quantification, isotope-labeled probes, metagenomic or metatranscriptomic measurements, and reaction-specific kinetic models could improve mechanistic insights. Nonetheless, data resulting from this assay is well-suited to inform microbiome-competent PBK models and other computational approaches that require quantitative parameters for gut microbial formation or depletion of bioactive chemicals. Moreover, biotransformation profiling contributes a unique level of functional characterization of complex microbial communities in the context of assessing microbiota-mediated exposure effects, bioactivation, and interindividual susceptibility to diet, drug, and xenobiotic exposures.

## Acknowledgements

ETH postdoctoral fellowship supported J.F. The European Partnership for the Assessment of Risks from Chemicals has received funding from the European Union’s Horizon Europe research and innovation program under Grant Agreement No 101057014 and has received co-funding of the authors’ institutions. Views and opinions expressed are, however, those of the author(s) only and do not necessarily reflect those of the European Union or the Health and Digital Executive Agency. Neither the European Union nor the granting authority can be held responsible for them.

## Data accessibility

Raw LC-MS/MS data files can be accessed from MetaboLights (https://www.ebi.ac.uk/metabolights/) under accession number MTBLS15096. Raw 16S rRNA gene amplicon sequencing data have been deposited in the European Nucleotide Archive (https://www.ebi.ac.uk/ena/browser/home) under project accession PRJEB115360. The analysis code repository is archived on Zenodo: https://doi.org/10.5281/zenodo.21528509

## Supporting Figures

**Figure S1.**
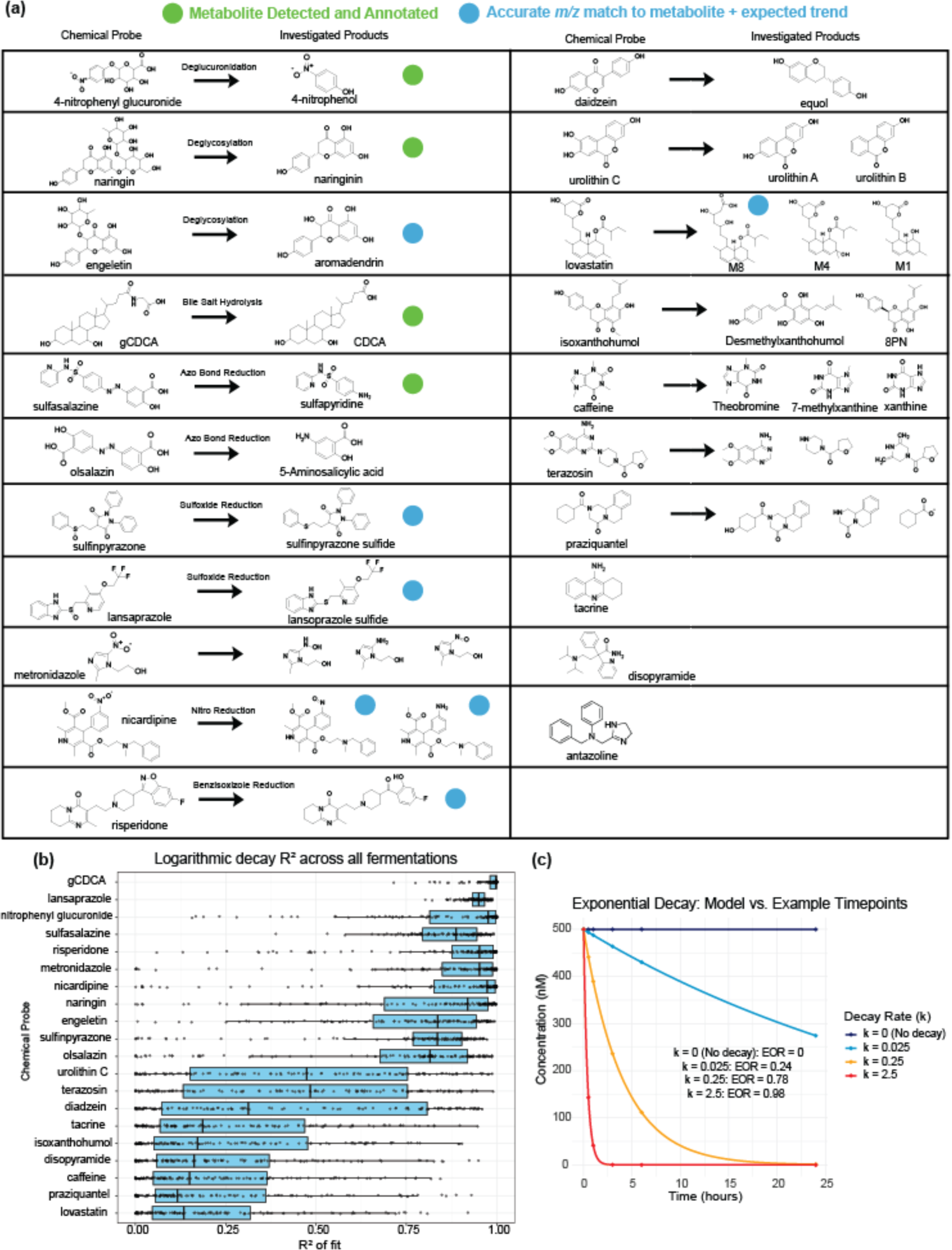
**A**. Green circles indicate metabolites that were annotated (*m/z* + MS/MS spectral match to reference library), blue circles indicate putatively assigned products based on accurate m/z and expected temporal trend (increases over time) and is present in samples with chemical probes but not probe-free control fermentations. The non-exhaustive selection of metabolites investigated were chosen by manual literature searches including the keywords “microbial metabolism” and the associated chemical name(s). **B.** R-squared values from logarithmic decay models fit for each fermentation and each probe separately. Probes are ordered from top to bottom as highest to lowest average R-squared value. **C.** Example dataset generated to compare logarithmic decay rate constants k and area under the curve based EOR metric for four different k values.

**Figure S2.**
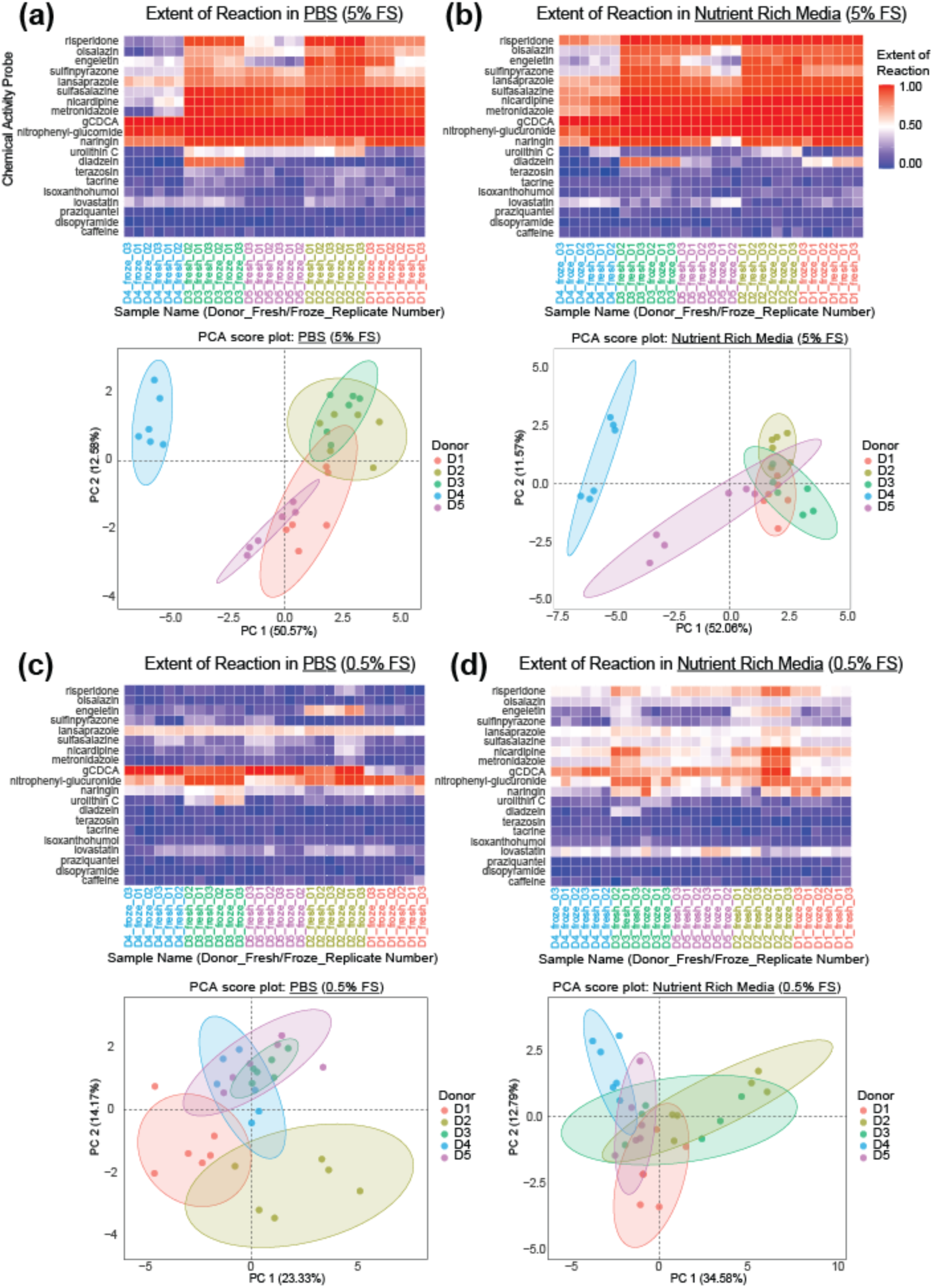
EOR value heatmaps and corresponding PCA score plots for four primary growth condition groups. PCA was performed on probe EOR values within each condition. Conditions tested are **A.** PBS media, 5% FS **B.** Nutrient rich media 5% FS **C.** PBS media, 0.5% FS and **D.** Nutrient media, 0.5% FS. Samples and probes were hierarchically ordered using Ward.D2, and order from PBS/5% fecal slurry condition was retained across panels for visual comparison.

**Figure S3.**
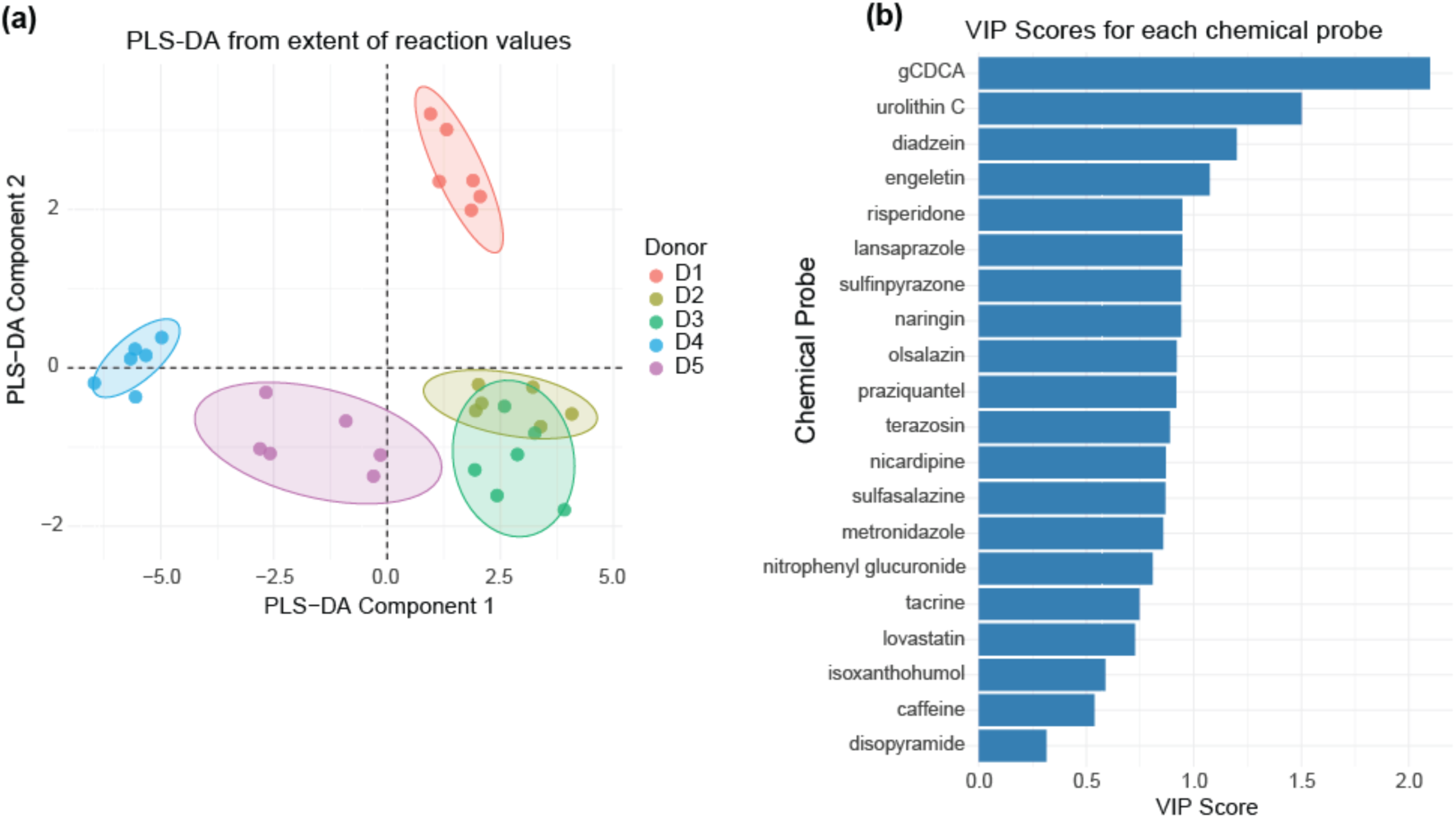
PLS-DA was performed on the hybrid EOR matrix, in which each probe’s EOR values were taken from the growth condition that produced the greatest inter-sample standard deviation for that probe. **A.** The score plot shows donor separation based on hybrid EOR profiles, and **B.** the VIP plot ranks chemical activity probes by their contribution to the PLS-DA model. VIP scores and PLS-DA calculations used “ropls” R package.

**Figure S4.**
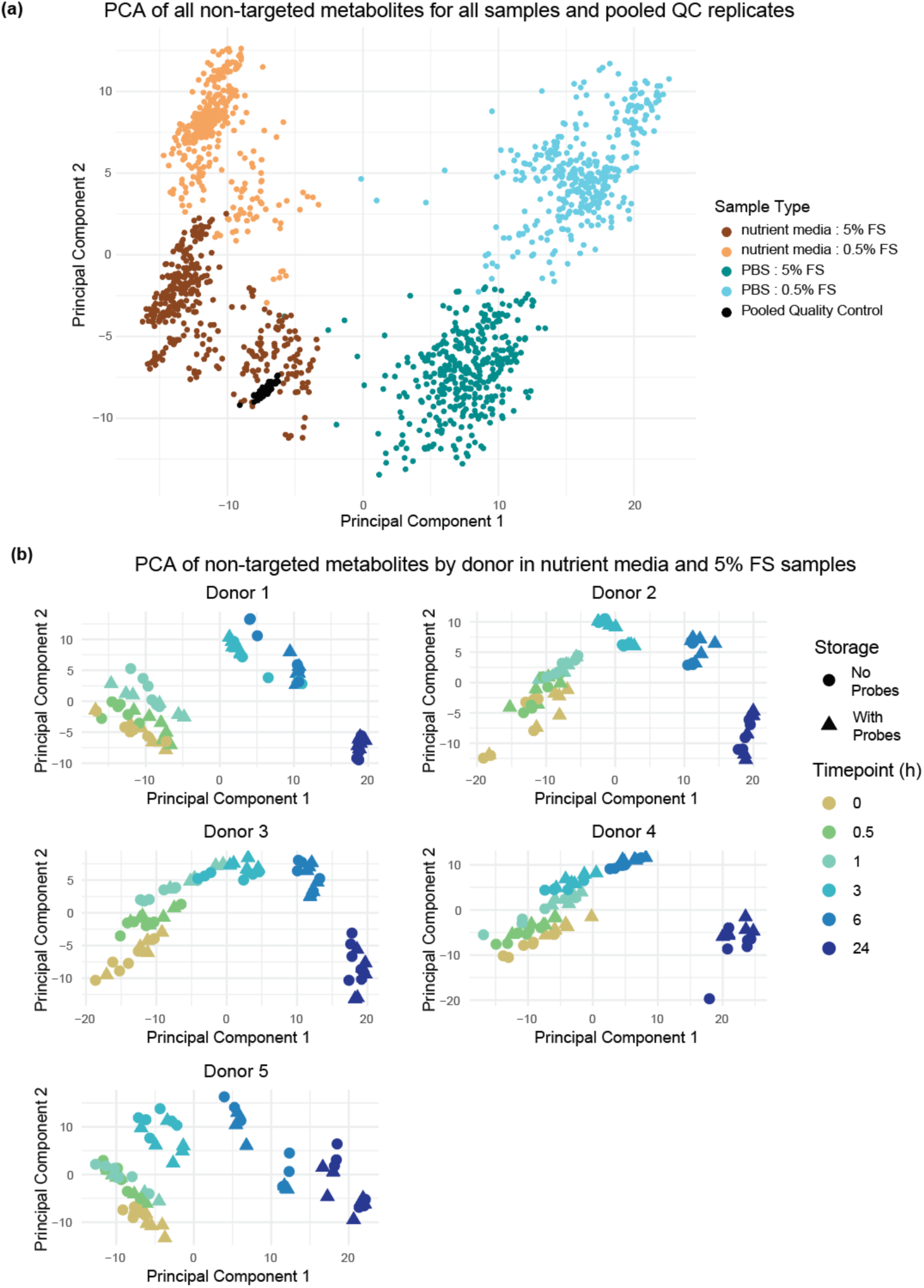
**A.** PCA of all samples and pooled quality control replicates using all annotated metabolites that were detected in at least 80 % of samples. **B.**PCA of metabolome data from each donor individually visualizes difference between samples with chemical probes added, or no probes added. Metabolome data from fermentations with nutrient media and high (5 %) FS inoculation growth conditions were plotted. Circle points indicate samples with no chemical probes added, while triangle points indicate samples with chemical probes added. Metabolite features with more than 20 % missing values were removed, as well as the chemical probe signals, leaving 312 metabolites included in this analysis. Missing values were imputed to 1/10 of the minimum value for each feature. Metabolite data was log10 normalized and scaled to unit variance.

**Figure S5.**
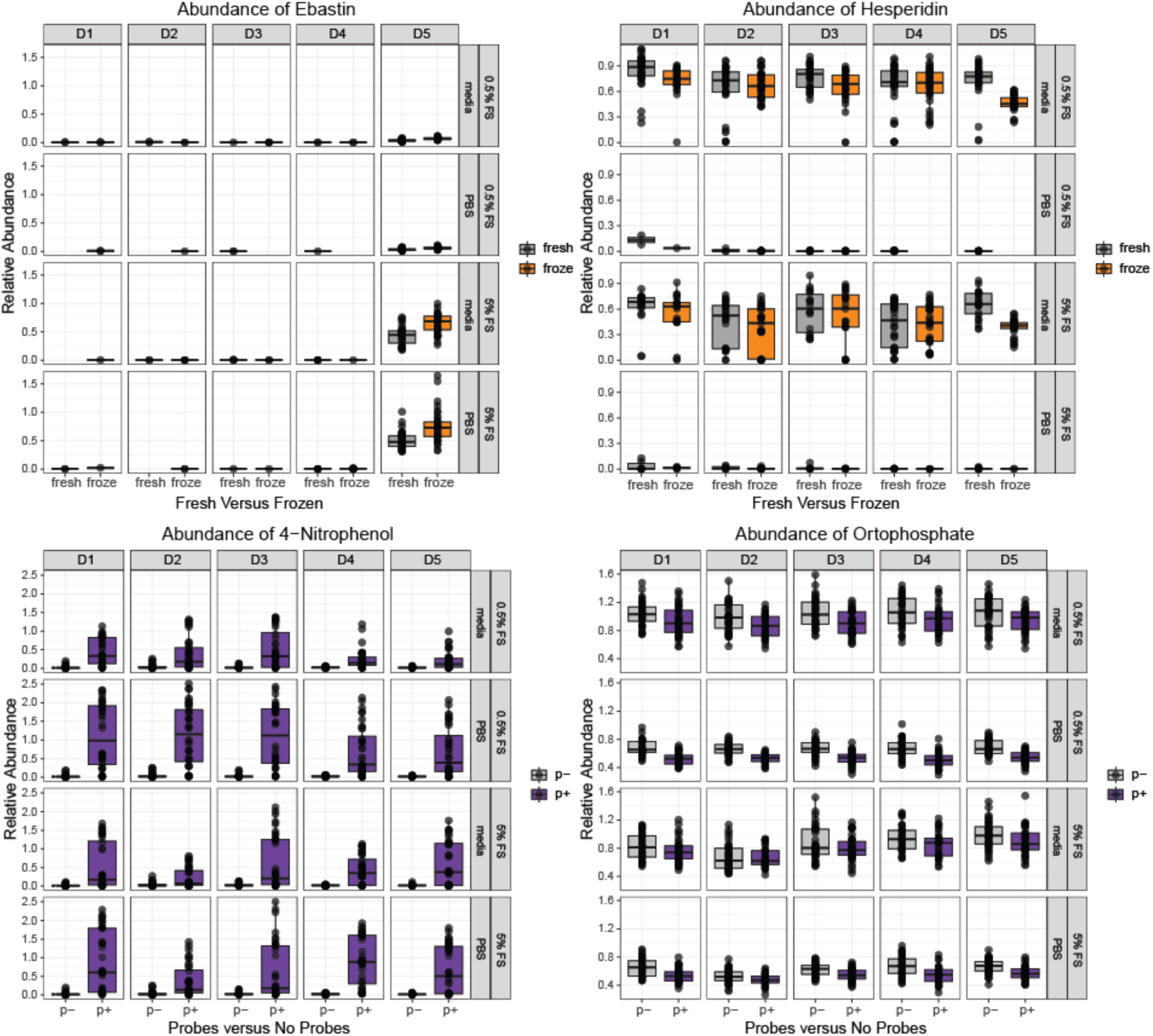
Boxplot of selected metabolites significantly changed based on treatments from non-targeted metabolomics analysis. Relative abundance represents untargeted metabolite peak height normalized within each sample to the sulfamethoxazole internal standard. Plots are faceted by donor, media/PBS background, and fecal slurry dilution. Points represent individual samples, and boxplots summarize distributions by fresh/frozen status or chemical-probe exposure, depending on the metabolite panel.

**Figure S6.**
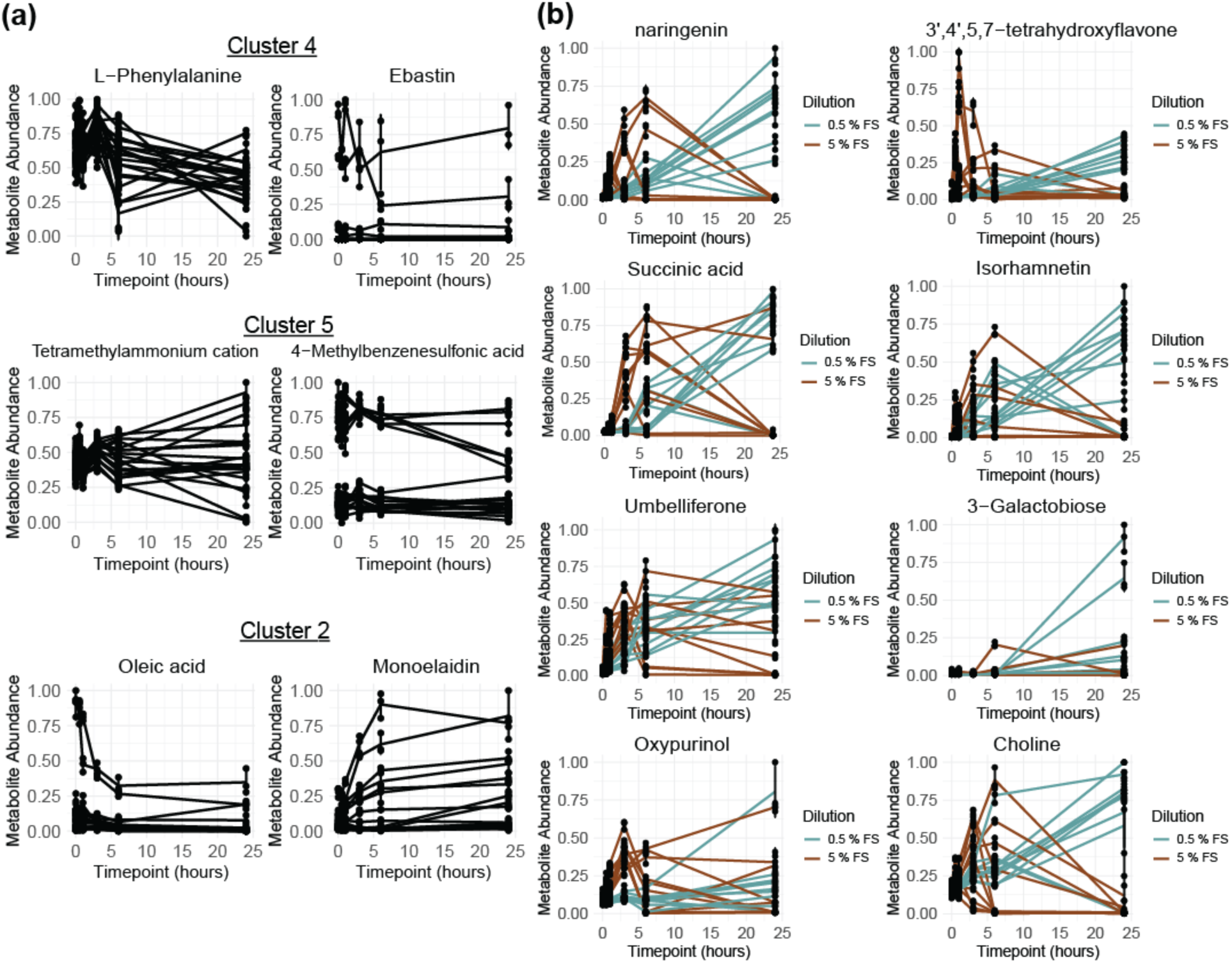
metabolite abundances over time plotted with all samples from media. Each point is an individual fermentation well from technical triplicates, and error bars are +/- 1 standard deviation (n=3). **A.** Representative metabolites chose from the indicated clusters. Profiles from 0 to 24 hours of fermentation displayed for samples containing probes and grown in media. **B.** Additional metabolites from cluster 6 are plotted differentiated by dilution level with 5 % FS inoculation in brown and 0.5 % FS inoculation in blue.

**Figure S7.**
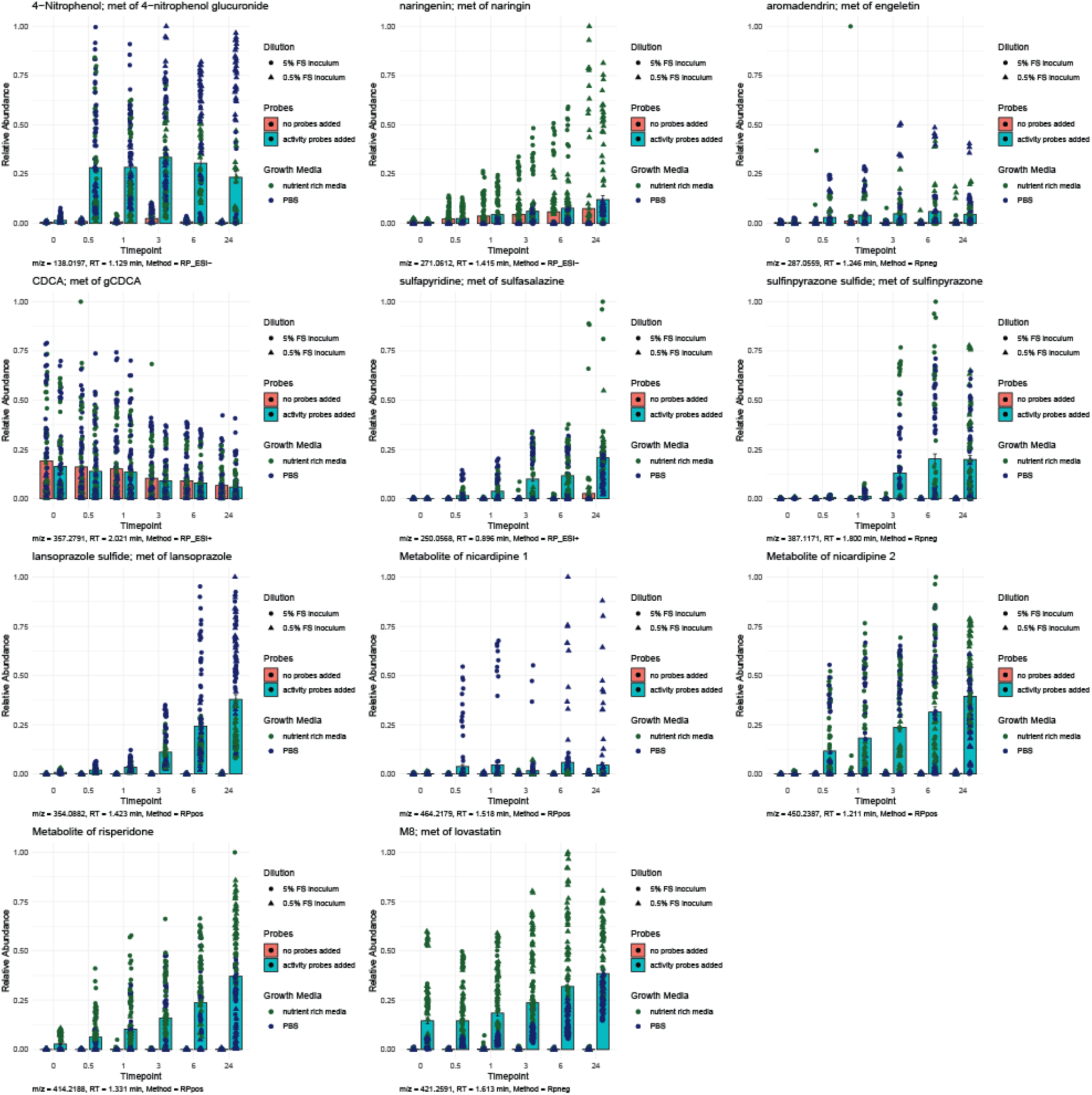
Abundance of metabolites detected in non-targeted LC-MS dataset. Each point represents one sample. Sampling timepoint in hours is presented along the x-axis, and normalized abundance (peak height scaled to 0-1) is plotted on the y-axis. The color of point describes whether the fermentation conditions were nutrient rich media (green), or nutrient scarce PBS (blue). Red or blue colored bars indicate whether chemical probes were added (blue) or not added (red) to the fermentation. Triangle points indicate 0.5% FS inoculation, while 5% inoculation samples are circles. The *m/z* of each LC-MS feature, retention time (RT), and whether it was detected in ESI+ (RPpos), or ESI- (RPneg) mode are indicated below each plot. Labels indicate the chemical name or identifier, and indicate that they are a metabolite (met) of the respective chemical probe.

**Figure S8.**
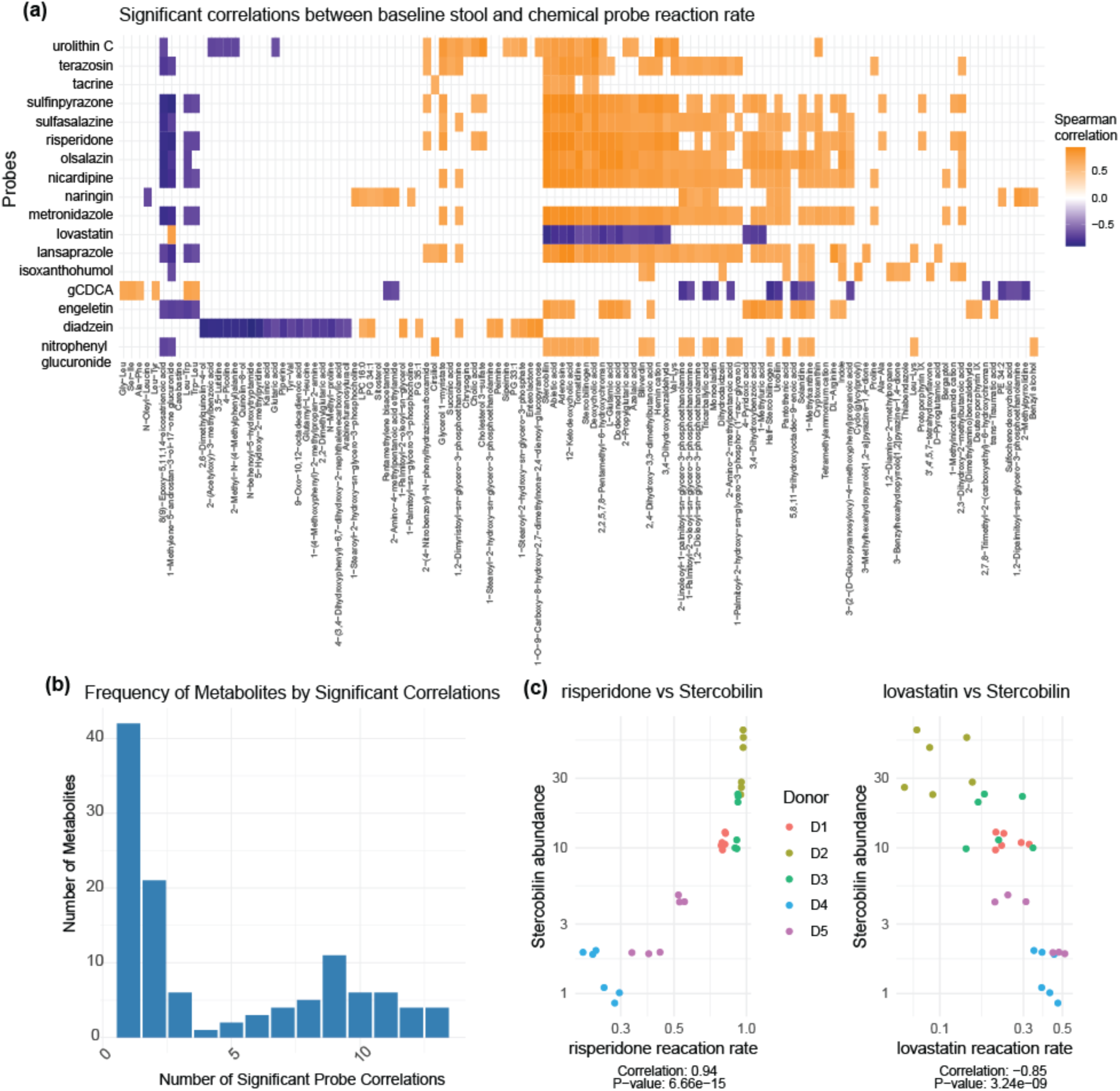
Chemical probe EOR values from growth conditions resulting in highest EOR variation were correlated to nontargeted metabolite abundances at timepoint 0 from PBS media and 5 % FS inoculation samples. In total 238 metabolites statistically significantly correlated (*p*<0.05, BH correction for n=6820 comparisons) to at least one chemical probe. To focus on only the strongest correlations, a more stringent filter of *p* less than 0.001 was applied resulting in 115 metabolites with at least one significant correlation to a chemical probe and visualized. **A**. Nontargeted metabolites are sorted by hierarchical clustering using method of ward.D. Panels B and C are histograms of how many non-targeted metabolites were significantly correlated to single probes highlighting the large number of metabolites that are correlated to multiple probes.

